# Librator, a platform for optimized sequence editing, design, and expression of influenza virus proteins

**DOI:** 10.1101/2021.04.29.441999

**Authors:** Lei Li, Olivia Stovicek, Jenna J. Guthmiller, Siriruk Changrob, Yanbin Fu, Haley L. Dugan, Christopher T. Stamper, Nai-Ying Zheng, Min Huang, Patrick C. Wilson

## Abstract

Artificial mutagenesis and chimeric/mosaic protein engineering have laid the foundation for antigenic characterization^1^ and universal vaccine design^2–4^ for influenza viruses. However, many methods used for influenza research and vaccine development require sequence editing and protein expression, limiting their applicability and the progress of related research to specialists. Rapid tools allowing even novice influenza researchers to properly analyze and visualize influenza protein sequences with accurate nomenclature are needed to expand the research field. To address this need, we developed Librator, a system for analyzing and designing protein sequences of influenza virus Hemagglutinin (HA) and Neuraminidase (NA). With Librator’s graphical user interface (GUI) and built-in sequence editing functions, biologists can easily analyze influenza sequences and phylogenies, automatically port sequences to visualize structures, then readily mutate target residues and design sequences for antigen probes and chimeric/mosaic proteins efficiently and accurately. This system provides optimized fragment design for Gibson Assembly^5^ of HA and NA expression constructs based on peptide conservation of all historical HA and NA sequences, ensuring fragments are reusable and compatible, allowing for significant reagent savings. Use of Librator will significantly facilitate influenza research and vaccine antigen design.

## Main

Influenza is considered to be the next major threat for a devastating pandemic. Vaccination has been proven an effective approach to prevent infection and global spreading of influenza viruses^6^. However, due to frequent mutations in the influenza virus genome, and particularly alterations to the HA and NA surface proteins, influenza vaccine protection is short-lived or mostly ineffective when mismatches have been observed for several flu seasons^7, 8^. In this context, developing universal influenza vaccine candidates that induce broadly reactive immunity and particularly antibodies against conserved epitopes of influenza virus surface proteins is an important direction of research^4, 9–12^. Use of artificial mutagenesis and design of antigen probes and chimeric/mosaic proteins are crucial steps in vaccine development and related research. However, current workflow for these tasks faces several major challenges that make it expensive, fallible and time-consuming. First, there are multiple residue numbering systems for HA protein sequences that have been commonly used in the literature, protocols and the research community^13, 14^. They are Coding Sequence (CDS) position, crystal structure-based H1/H3 numbering^15–17^, and Burke and Smith HA numbering. For a given sequence, CDS position usually counts from the start of the CDS, methionine, and therefore can cover all amino acids; structure-based H1/H3 numbering and Burke and Smith HA numbering determine residue numbers according to template mapping using different templates (see methods section for details). Thus, biologists must put significant effort into identifying the correct residues to avoid errors. Moreover, nucleotide and amino acid sequences are difficult to read, and inefficient and fallible to edit manually. Further, there are no comprehensive tools to develop individual influenza sequence databases and readily compare varied influenza sequences and phylogenies, or to immediately port annotated sequences for visualization of color-coded antigenic-regions, mutations, or epitopes on representative HA and NA structures using structure analysis software such as PyMol and UCSF Chimera. Finally, efficient and automated cloning of HA and NA protein variants to scale can become expensive and is error-prone if done by hand. Gibson Assembly can assemble multiple linear DNA fragments for protein cloning and expression and has been extensively used in molecular biology, and is superior among all assembly methods because of the enormous savings of time and human labor with its easy one-tube reactions. Automated Gibson fragment prediction and databasing of fragments conserved between similar HA and NA protein expression constructs would allow accurate and cost-effective production of variant influenza protein libraries for many applications. In conclusion, these broad challenges in current influenza immunology/virology studies limit the efficiency and breadth of research. Improving the accuracy and economy of these processes will significantly expand related studies both in seasoned influenza laboratories and for novice laboratories interested in applying innovative approaches to the study of influenza.

Here, we present a computational tool called “Librator” for influenza HA and NA sequence analysis, editing, and cost-effective cloning and vector design for HA and NA protein expression. Librator is an integrated graphical processing platform for influenza sequences. Librator seamlessly connects nucleotide sequences (from public sequence databases) and lab work (e.g. Gibson cloning, protein expression), and it contains a variety of functions to facilitate management, analysis, editing and accessing influenza sequences, to improve the efficiency of sequence design and expression (Figure 1A). This software entrusts all error-prone sequence editing and data processing tasks to the background algorithm, so that users are able to design their sequences with a few clicks on the GUI. With Librator users can complete all sequence design-related operations graphically in an integrated system, avoiding difficult-to-read raw sequences or switching between different software applications.

**Figure 1.**
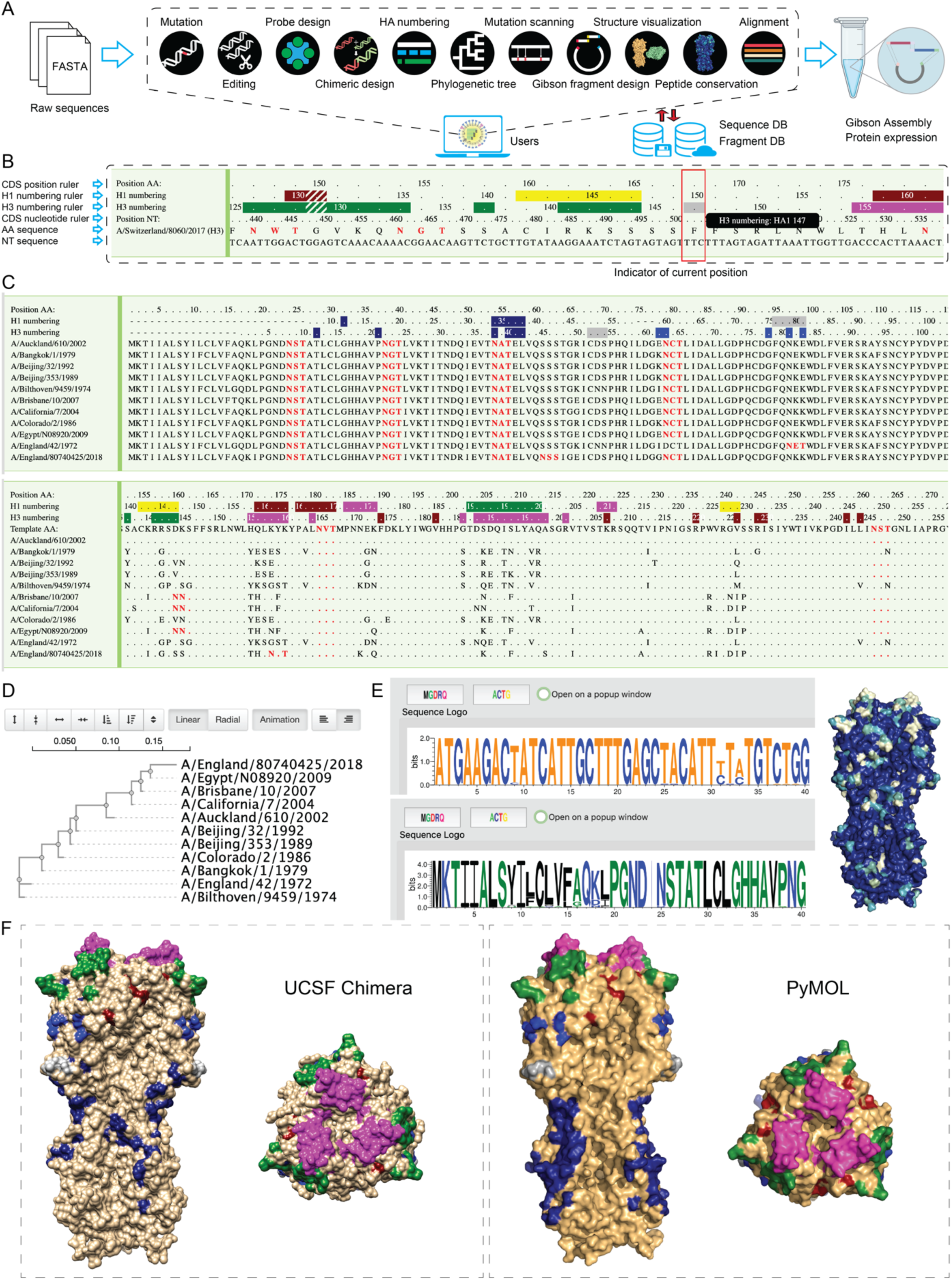
Librator enables efficient analysis of HA and NA influenza virus protein sequences with a variety of functions and a graphical user interface. (**A**) Librator seamlessly connects nucleotide sequences from public databases and lab work, providing a variety of functions for sequence editing and design. (**B**) Librator’s HA numbering aligner is integrated in a graphical viewer. Three numbering rulers—a CDS position ruler, H1 numbering ruler and H3 numbering ruler—indicate position information for selected residues. (**C**) Multiple sequence alignment viewer. Original sequence mode is displayed on the top, and template mode is displayed on the bottom. (**D**) Phylogenetic analysis function and tree viewer. Users are allowed to generate phylogenetic trees using either nucleotide sequences or peptide sequences. (**E**) Librator allows users to assess nucleotide conservation and peptide conservation. Librator also can visualize the peptide conservation on HA 3D structures with the help of PyMOL and USCF Chimera. (**F**) Librator allows users to visualize peptides on 3D structures of HA proteins with color-annotated amino acids at all antigenic regions and user-defined sequence labels with the help of PyMOL and USCF Chimera.

Multiple common functions were built-in to Librator to help users analyze influenza sequences more efficiently. First, an HA numbering aligner was integrated into Librator’s sequence viewer and editor. This function aligns given HA sequences to crystal structures of a classic H1 (PDB ID: 4JTV) and a classic H3 template (PDB ID: 4HMG) to identify corresponding H1/H3 numberings for each residue. Known common antigenic sites and epitopes are automatically labelled based on this numbering; for example antibody binding sites (ABS), receptor binding sites (RBS), and the H1 (Ca1, Ca2, Cb, Sa, Sb, Stalk) and H3 (A, B, C, D, E Stalk) antigenic sites are color-coded on H1/H3 numbering rulers (Figure 1B). Notably, epitope definitions in Librator are highly customizable, allowing users to annotate HA sequences according to their specific research interest and focus (Figure S1A). Since glycosylation on the HA protein was reported to have an important impact on antigenic drift^18, 19^, an “N-X-S/T” pattern that indicates potential N-linked glycosylation sites are also highlighted. This viewer is also capable of displaying fully annotated multiple sequence alignments with two informative modes, original sequence mode and template mode, enabling convenient investigation of evolutionary sequence patterns and mutations (Figure 1C). For example, biologists characterizing escape mutant sequences induced by selective pressure with antibodies or sera can align the HA or NA sequences and immediately visualize and export graphics of regions that were mutated. To help users quickly infer phylogenetic relationships among a group of sequences, Librator also allows users to generate and visualize maximal likelihood trees from either nucleotide sequences or peptide sequences (Figure 1D). Furthermore, powered by WebLogo, Librator allows users to access nucleotide and peptide conservation among groups of sequences^20^ (Figure 1E). By automatically porting amino acid sequences and labelling instructions to PyMOL^21^ or UCSF Chimera^22^, Librator allows users to visualize peptides on 3D structures of HA proteins with color-annotated amino acids according to either peptide conservation score (Figure 1E) or all antigenic regions and user-defined sequence labels (Figure 1F). Librator uses an H1 structure (PDB ID: 4JTV) for visualization of all Group 1 HA structures and a H3 structure (PDB ID: 4HMG) for visualization of all Group 2 HA structures^23–25^. For example, with a single button-click, users can immediately evaluate whether an escape mutation is predicted to alter a surface amino acid or occurs deeper in the structure potentially driving conformational changes. In addition, a function was also developed that allows users to identify potential key residues between two groups of sequences by ranking residues by their amino acid difference. For example, by comparing pre-1994 and post-1994 human H1N1 seasonal viruses, Librator highlighted the importance of a deletion “Δ130,” which has been validated by experiments^26^, by a high ranking score. This tool helps users zero in on important sequence elements driving influenza evolution. Lastly, an independent viewer for users to easily access the Burke and Smith HA numbering scheme proposed by Burke et al. was implemented since it has also been commonly used in the Influenza community^13^(Figure S1B). It should be noted, however, that all functions in Librator, including the alignment viewer, sequence editing, and sequence designing, were based on structure-based numbering systems.

To improve the efficiency and accuracy of mutagenesis and sequence editing, we developed multiple functions to help users to design and edit their Influenza sequences. With the help of the HA numbering aligner, users can easily locate target residues and mutate them by simply typing a mutation code using whichever numbering system they prefer. For example, for an H3 sequence (A/England/80740425/2018), typing “Y177M” in CDS position input will mutate the 177^th^ residue of the CDS from Tyrosine (Y) to Methionine (M). This is equivalent to typing “Y164M” in H1 numbering HA1 input or typing “Y161M” in H3 numbering HA1 input (Figure 2A). By translating between the various numbering schema, Librator avoids confusion and mistakes that are common in analyzing influenza sequence data. For NA sequences, only CDS position input is available since it is the only numbering system for NA sequences. To avoid mistakes, Librator validates the original amino acid in the mutation code to make sure it matches the amino acid in the raw sequence in the numbering system used. Expression of influenza HA soluble proteins for experimental purposes is an important tool for characterizing influenza immunity or monoclonal antibody specificity. Building on this mutagenesis function, we also developed a function to design HA expression constructs for most HA subtypes (H1–H15, see methods section for details) with one click that replaces the flexible linker and transmembrane region with a stabilizing Trimerization domain, an Avitag for mono-biotinylation, and a histidine six-mer (H6) sequence for nickel-based purification (Figure 2B)^27^. Using the “probe option” of this function also introduces a “Y98F” mutation (H3 numbering) that reduces binding to sialic acid for probes to be used in cellular assays such as for flow cytometric sorting of HA-specific B cells^28^ or Libra-seq^29^.

**Figure 2.**
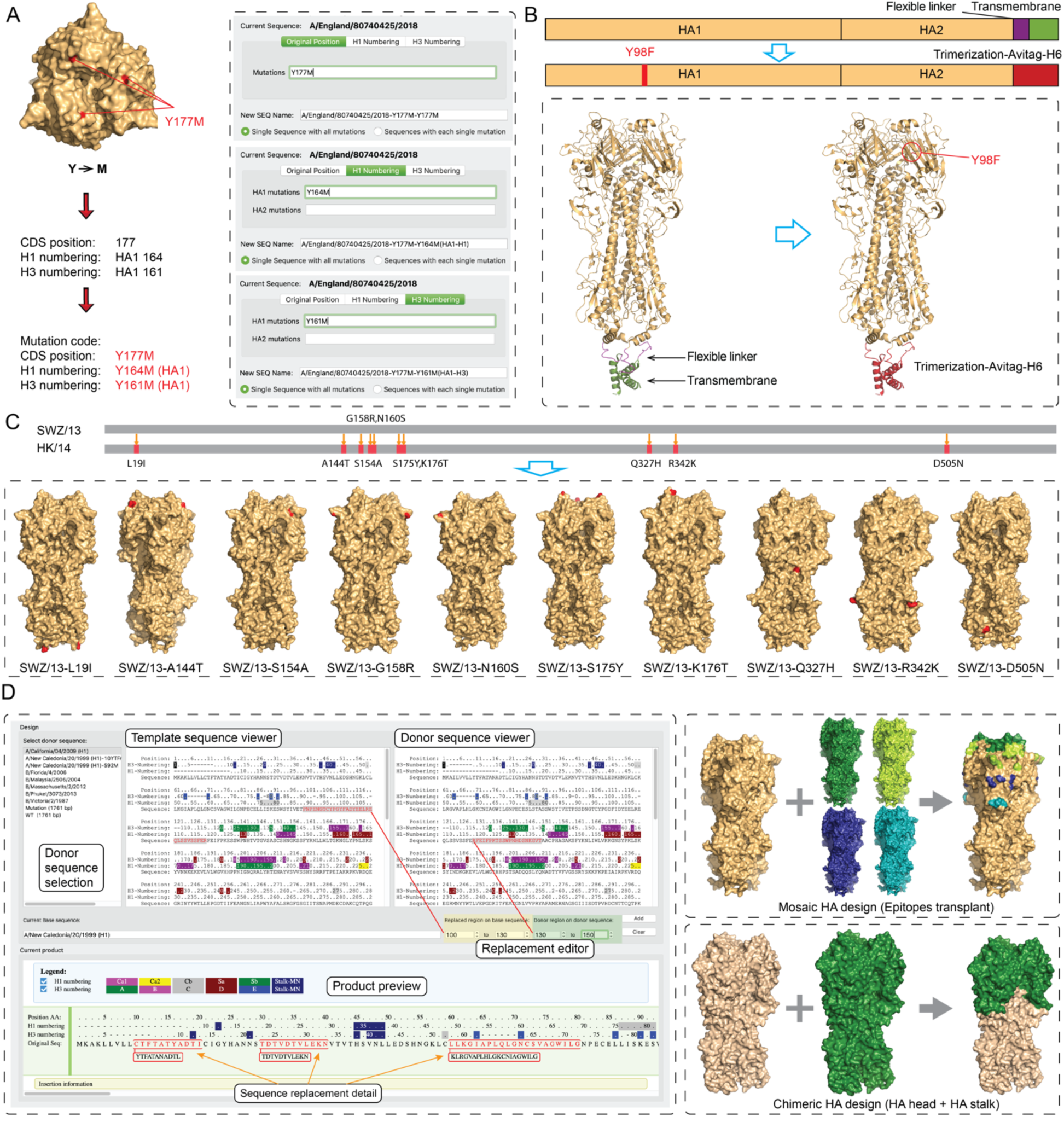
Librator enables efficient design of HA and NA influenza virus proteins. (**A**) Demonstration of mutating a residue on an HA sequence using three numbering systems in Librator. (**B**) Making an antigen probe for a given HA sequence. Librator designs antigen probes for given HA sequences by generating the mutation Y98F (H3 numbering) and replacing the flexible linker and transmembrane region with a Trimerization-Avitag-H6 sequence. This process is demonstrated using an HA structure of A/duck/Alberta/35/76 (H1N1, PDB ID: 6HJR). (**C**) Scanning all amino acid differences between two antigenically distinct sequences (A/Switzerland/9715293/2013 [SWZ/13] and A/HongKong/4801/2014 [HK/14]) with Librator generates a series of sequences, each with a single mutation, to identify key residues of the antigenic drift. (**D**) Designing chimeric sequences using Librator. Users can replace regions on the target sequence with regions from multiple donor sequences. Details of the product can be previewed on a graphical viewer. This function is designed to transplant epitopes from one sequence (or multiple sequences) to another or to combine the HA1 (HA head) from one sequence and HA2 (HA stalk) from another.

In addition to the mutagenesis function, Librator also provides several sequence editing modes. For example, the Cocktail mode allows users to compare a donor sequence to a template sequence and scan all amino acid differences between them. Then Librator will automatically generate multiple sequences based on the template sequence with each identified mutation or their combinations (Figure 2C). This function improves the efficiency of identifying key residues between antigenically or functionally distinct viruses. Users can instantaneously generate a library of point mutant variants for expression to, for example, identify which amino acids differing between two HA molecules are important for the binding of a monoclonal antibody or drive differential function of the compared HA molecules, such as host-species tropism. With these functions, users can generate demanded mutations in batches in minutes, compared with manual generation of mutated sequences that usually takes at least several hours.

We also implemented a sequence designer in Librator to facilitate complicated sequence design. Current influenza vaccine-antigen design efforts aim to retarget immunity away from some epitopes and focused on others through the production of chimeric and mosaic HA and NA proteins. Compared to individual mutagenesis, chimeric and mosaic sequence design usually requires mutating multiple regions (groups of residues) or even splicing sequences together from multiple influenza strains. Large numbers of mutations and complex design make manual design of chimeric/mosaic HA proteins difficult and prone to error. To overcome this challenge, Librator includes an interactive GUI to enable easy and efficient design of chimeric and mosaic proteins (Figure 2D). Using the graphical sequence viewer, users can easily specify and highlight regions to be replaced on a template sequence and regions to be inserted from a donor sequence. A dedicated viewer displays the current product with information about all replacements. After users review and confirm the current product in the product viewer, Librator will generate a new record of the user-designed product, with nucleotide sequence, subtype (same as template) and mutated residues. Using Librator, biologists can easily design complicated chimeric and mosaic HA/NA sequences with extensive mutations or replacement of entire epitopes or regions. Use of Librator in our lab has enormously improved the efficiency and accuracy of sequence design.

Librator’s cloning functions also maximizes the economy, practicality, and accuracy of synthesizing nucleotide sequences for expression by Gibson cloning using a recipe-based generator. For this Librator capitalizes on the fact that Gibson cloning uses sequence homology of a short overlap/joint region (usually 20–25bp) between neighboring fragments and also that most HA and NA sequences of a type have highly conserved and homologous regions interspersed with the variable sequence elements. Natural mutations in these proteins are enriched in only a few highly variable regions (e.g. epitopes, antibody binding sites) (Figure 3A). The cloning algorithm of Librator optimizes fragment design for HA and NA sequences to maximize the reusability of gene fragments. Librator typically produces HA as four fragments (user customizable) or NA as three fragments and databases all previous fragments generated by a lab so that new HA molecules differing in only one fragment can be synthesized by replacing only the single fragment based on an automatically generated recipe specifying the existing fragments in the laboratories inventory and the new sequence to be synthesized (Figure 3B). For example, an escape mutant HA of a particular strain may contain only several amino acid changes within a single antigenic site in one fragment of the construct. If the original variant was designed by Librator and expressed in the lab, the escape variant can now be synthesized at only 1/4^th^ the cost. This function become particularly cost-effective when libraries of point-mutants are generated. For this, Librator identifies potential overlapping regions by locating highly conserved regions based on peptide conservation of all historical HA and NA sequences. These regions are then used to define fragments on a template sequence for each subtype or group of subtypes, ensuring that end compatibility of fragments is unaffected by sporadic mutations, insertions or deletions. In Librator, all query sequences are aligned to the appropriate template sequence to ensure fragments from different batches are subject to the same design, guaranteeing their reusability. Users can clone and express their HA and NA sequences for a reduced cost by reusing fragments in their inventory (Figure 3C). The more sequences users clone, the more comprehensive a fragment inventory they will amass, enabling more fragment reuse and reagent saving. This is extremely beneficial for labs that are investing continuing efforts and resources into influenza research.

**Figure 3.**
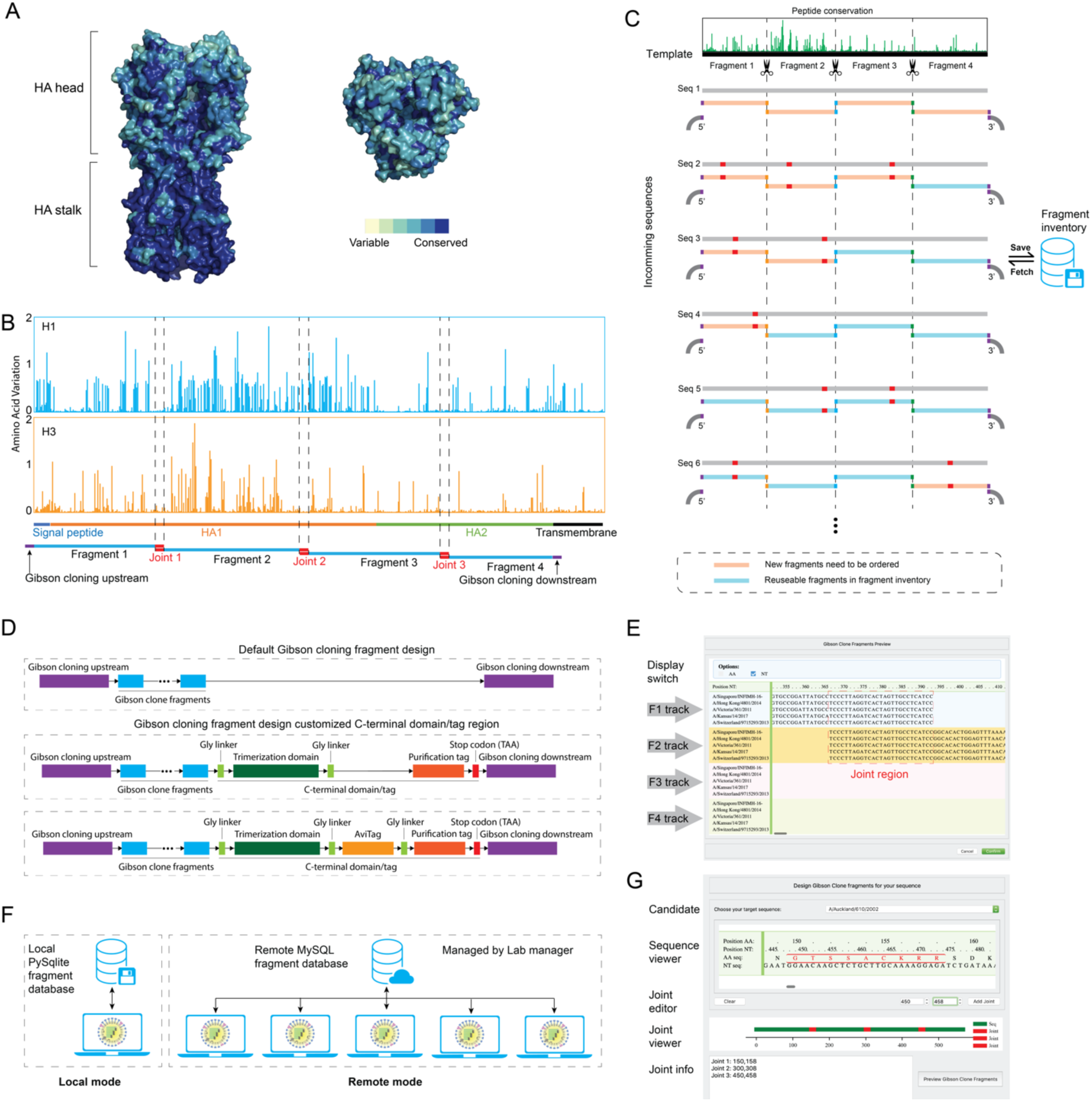
Librator helps users to save reagent cost by designing optimized Gibson Clone fragments for HA and NA sequences. (**A**) Natural mutations on HA protein are enriched in a few highly variable regions. Peptide conservation was visualized on a H1 protein structure (A/California/04/2009 H1N1, PDB ID: 4JTV). Peptide conservation was calculated from HAs of 58 representative H1N1 viruses from 1918 – 2018. (**B**) Illustration of fragment designs for a group 1 HA (based on a H1 template) and a group 2 HA (based on a H3 template) in Librator. Joint regions were determined by locating highly conserved regions on H1/H3 peptides and balancing the length of each fragment. (**C**) Librator determines overlapping regions based on peptide conservation of all historical HA and NA sequences, and then defines fragment design on a template sequence for each subtype. For each given sequence, Librator aligns it to its template sequence to maximize the reusability and compatibility of gene fragments. All fragments are saved in a fragment inventory for further inquiry. Librator aims to save reagent cost by reusing gene fragments. (**D**) Three modes of customizing the C-terminal domain/tag region for the Gibson cloning downstream end. Beside the default mode that directly links the last fragment and the Gibson cloning downstream sequence, we also designed a customizable C-terminal domain/tag region for HA proteins: Trimerization domain + Purification tag (e.g. 6xHisTag) or Trimerization domain + AviTag + Purification tag. (**E**) Graphical viewer of fragments for users to preview their products. (**F**) Librator users can communicate with a local fragment database or a remote fragment database managed by their lab manager. The remote mode enables better data access and lab reagent stock management. (**G**) Customized fragment design function for any given sequence. This function allows users to add at most 12 joint regions in their sequences and split their sequences into a few fragments for Gibson Assembly. This function was designed for non-influenza sequences or novel research in which reusability is not a priority.

According to the evolutionary history of influenza HA subtypes, we designed uniform fragments on the basis of a classic H1 sequence (A/California/7/2009, H1N1) and a classic H3 sequence (A/Aichi/2/1968, H3N2). We aligned all group 1 HAs (H1, H2, H5, H6, H8, H9, H11, H12, H13, H16, H17, and H18) to the H1 template and all group 2 HAs (H3, H4, H7, H10, H14, and H15) to the H3 template for fragment design (Table S1,S2, Figure S2). For NAs, we designed uniform fragments for each subtype by aligning each of the NA sequences to the template of their respective subtypes (Table S1, Figure S3). This template-mapping-based fragment design ensures that all fragments are standardized and not affected by either different batches or sporadic insertion/deletion events (e.g. a deletion Δ130 between pre-1994 and post-1994 human seasonal H1N1, or insertions in the cleavage site of high pathogenic avian H5 and H7)^26, 30, 31^. We applied this system to several applications to validate its effectiveness and compatibility. Lab practices demonstrated that this tool could help to clone and express proteins at a reduced cost. For example, reagent cost was reduced by 54% when expressing proteins with single mutations to investigate the key residues of the antigenic drift between A/HongKong/4801/2014 (H3N2) and A/Switzerland/9715293/2013 (H3N2) influenza viruses (Supplemental Data S1). Even in an extreme case of expressing HAs of 39 representative H3N2 viruses from 1968 to 2018, using Librator design only increased the reagent cost by 4% while generating many reusable gene fragments for future projects (Supplemental Data S2).

With the effectiveness of this method verified by lab practice, we further developed several supporting functions to enable efficient workflow and a smooth user experience. We enabled users to customize the Gibson upstream connector and downstream connector to fit more vectors. Furthermore, we also designed a customizable C-terminal domain/tag region for HA proteins: Trimerization domain + Purification tag (e.g. 6xHisTag) or Trimerization domain+ AviTag + Purification tag (sequences are user customizable) for better end compatibility (Figure 3D). To reduce the risk of error, we designed an interactive GUI on which users can preview their designed fragments before generating all products (Figure 3E). All generated fragments are archived in an SQL-driven database for better data access and management. To facilitate lab reagent stock management, Librator also allows multiple users to connect to a remote MySQL fragment database (Figure 3F). Once fragments are generated by users, Librator searches the current fragment inventory, then generates a list of reusable fragments already in inventory, novel fragments that need to be ordered and recipes for all sequences. An Excel file containing fragment names and sequences in the format of a 96-well plate is also generated and can be sent to a DNA synthesis company directly. FASTA format files that contain the fragments of each sequence are generated as well, enabling users to validate their compatibility using sequence analysis software. Lastly, we also developed a general fragment design feature that allows users to split any nucleotide sequence into a few customized fragments, most applicable when reusability is not a priority (Figure 3G). This feature will be helpful for novel or frontier research in particular, such as in designing Gibson cloning fragments for novel COVID-19 proteins.

In conclusion, we developed a variety of functions associated with interactive GUIs in Librator, aiming to improve research efficiency and liberate biologists from onerous and repetitive work so that they can focus on more productive aspects of influenza research. This feature greatly facilitates the work of users who are not familiar with command-line tools, as well as reducing the possibility of mistakes. Furthermore, by the help of two widely used structure visualization tools, Librator seamlessly links users linear HA sequences to 3D structures that are annotated by peptide conservation and known epitopes. This unique feature facilitates virologists, especially those who are not expertise in structural biology, to investigate their sequences and designs from structural aspect. We also provide tools for optimized Gibson clone fragment designs for HA and NA proteins of influenza viruses, enabling low-cost protein cloning and expression. This protocol liberates more scientific potentials for related research under limited budgets, expending the depth and breadth of related research. Looking to the future, Librator has much potential to be extended. In recent years, more and more studies have revealed epitopes on NA proteins and highlighted the importance of NA as a target of human antibodies^32–35^. Compared to HA, there is still a lack of knowledge of NA. In the near future, more and more studies will focus on NA and will be able to generate comprehensive profiles of epitopes on NA. Librator will be continuously updated with the latest research progress on NA. Furthermore, compared to influenza A, there is a lack of knowledge about influenza B, which also has an impact on public health and is also an important component of WHO-recommended influenza vaccine formulas. Improving support for influenza B is another future goal for Librator. Lastly, this template-based and standardized fragment design also has the potential to be extended to other viruses, such as human immunodeficiency virus (HIV) or hepatitis C virus (HCV) or coronaviruses. The modularized structure of this software is also ready for secondary development to be compatible with more biological contexts. With this in mind all source code is provided and we encourage updates and feedback and hope that Librator becomes a community-based tool and development effort.

## Methods

### Dataset

All the HA and NA sequences used in this study were downloaded from the NCBI FLU database (https://www.ncbi.nlm.nih.gov/genomes/FLU/)^36^ and GISAID database (https://www.gisaid.org/)^37^. H1 protein: 2243 seasonal H1 sequences and 31575 pdm09 sequences. H3 protein: 61798 sequences. NA protein: 28747 N1 sequences, 15194 N2 sequences, 1430 N3 sequences, 291 N4 sequences, 382 N5 sequences, 2420 N6 sequences, 1188 N7 sequences, 2446 N8 sequences and 2446 N9 sequences. All sequences are peptide sequences.

### Gibson Clone fragment design for HA and NA proteins

Gibson Clone fragments should be designed according to a uniform criterion that is unaffected by sporadic insertions/deletions in different strains, and all the joint regions of neighboring fragments should be located at the most conserved region. Furthermore, an optimized fragment design should also balance the reusability of each single fragment and the total number of fragments. The shorter a single fragment is, the less the probability of mutations will be, enabling higher reusability of each fragment; too short a fragment length will result in a larger number of fragments, however, which highly increases the total reagent cost.

To determine the optimized fragment design (including number of fragments and joint region location), we investigated amino acid variations of all residues of human H1, human H3 and NA (all hosts), and we quantified the amino acid variations by an amino acid variation entropy function^20^.

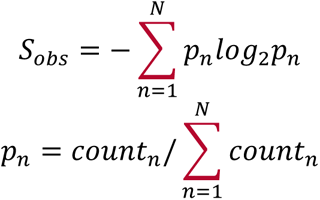

*S_obs_* denotes the entropy of the observed symbol. *p_n_* denotes the frequency of the *n*-th amino acid of this residue, *N* denotes the total number of all possible amino acids (N = 20), and *count_n_* denotes total number of the +-th amino acid of this residue.

By comprehensively considering commercial DNA fragment sizes and prices and distribution of the conserved regions in the HA/NA sequences, we proposed an optimized fragment design that divides HA into 4 fragments and NA into 3 fragments. Length of joint regions was set to 9 amino acids (27bp in nucleotides) because Gibson Clone Assembly requires at least 25bp joint region length. Joint regions of group 1 HA were set at 123–131, 264–272, and 403–411 (CDS position on a A/California/7/2009[H1N1] HA). Joint regions of group 2 HA were set at 123–131, 265–273, and 403–411 (CDS position on a A/Aichi/2/1968[H3N2] HA). Joint regions of NA were set at 131–139 and 292–301 (for each subtype, all positions are subject to CDS position on a representative template of this subtype). Joint regions and templates of all HA and NA subtypes are shown in Table S1. Furthermore, to maximize the compatibility of joint regions, Librator revised all nucleotide sequences of joint regions by translating them from peptide sequences using a dictionary in which each amino acid only has one corresponding codon.

### Pipeline design

To optimize the user experience, especially for biologists without a computer science background and not familiar with command-line tools, we developed a highly interactive GUI for Librator. Function calling, parameter setting and information display were integrated into one main interface with multiple tabs. All functions can be divided into two broad categories: basic function and advanced function. Basic function includes Input/output (I/O) operations and database (DB) operations: parameter setting, create new sequence DB, open existing sequence DB, import sequences and export sequences. Advanced function inGUIcludes sequence design/editing, fragment design, phylogenetic analysis and structure visualization: the specific functions are sequence information editing, HA numbering, mutation identification, antigen probe design, multiple sequence alignment, phylogenetic analysis, sequence editing, chimeric HA design, structure visualization and Gibson Clone fragments design (Figure S4).

### Multiple HA numbering schemes in Influenza research field

As discussed in the introduction section, there are three different numbering systems commonly used in the Influenza research field: 1) CDS position, 2) crystal-structure-based H1/H3 numbering, and 3) Burke and Smith HA numbering scheme.

Residue number on CDS is usually counted from the first amino acid of the CDS (Methionine). For a given sequence, CDS position can cover all residues of given sequence regardless of sporadic insertion and/or deletion. The crystal-structure-based H1/H3 numbering aligns given sequences against a classic H1/H3 template and assigns position numbers for all residues that can map to the template crystal structures. Thus, inserted residues and non-structural residues (e.g. signal peptides) will not be assigned a residue number because they cannot be aligned to the template crystal structures. Furthermore, numbers of residues in HA1 and HA2 are counted independently. The Burke and Smith HA numbering scheme proposed by Burke et al. aligns given sequences against 26 templates of different subtypes to determine the residue numbers. Different from the structure-based HA numbering scheme, the Burke and Smith HA numbering scheme is based on amino acid sequences without considering structural information, and it counts from the first amino acid of the CDS after signal peptide removal. This numbering scheme has been implemented by FLUDB (https://www.fludb.org/brc/haNumbering.spg?method=ShowCleanInputPage&decorator=influenza) recently. We compared three different HA numbering systems using H1 and H3 template sequences (Figure S5; Table S3, S4).

Because protein structures play an important role in antigen phenotypes, all functions in Librator, including alignment viewer, sequence editing and sequence designing were based on structure-based HA numbering systems. Users can only access the universal HA numbering scheme in the “Burke and Smith HA numbering” viewer.

### Antigen probe design

The antigen probe design function makes HA probes for a given HA sequence by generating a “Y98F” mutation (H3 numbering) and replacing the flexible linker and transmembrane region with Trimerization-Avitag-H6 sequence. Residue 98 under H3 numbering is located by the built-in HA numbering aligner system automatically. The transmembrane region is identified by aligning given sequences to an H3 template. This function is not available for most H16 HAs and all H17 and H18 HAs because these sequences are isolated from avian and bat sources, and their residue 98 under H3 numbering is already “F.”

### Identification of key residues between two groups of sequences

In this function, first we align all sequences from both groups together; then we investigate peptide differences between the two groups for every residue independently. For each residue, we convert amino acid composition of two groups into numerical amino acid vectors:

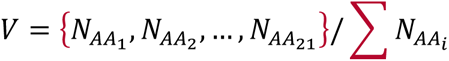

*AA_i_* denotes the 6-th amino acid of a total of 21 different amino acid options (20 amino acids + any symbol beside those 20 AAs, e.g. alignment gap or unclear amino acid X). *N_AAi_* denotes the total number of appearances of the *i*-th amino acid. Then we defined a score to represent the difference in amino acid composition between two groups on a specific residue:

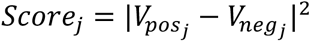

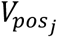 and 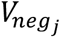 denote amino acid vectors of residue *j*. *Score_j_* denotes peptide difference of residue *j*.

Under this scoring system, score = 0 indicates no peptide difference on this residue between two groups. The higher the score is, the bigger the peptide difference will be. Then all residues will be ranked by the score from high to low to facilitate users’ further analysis. In summary, this function gives suggestions of key residues to narrow the candidate range and accelerate biological studies.

### Nomenclature of Gibson Clone fragments

Users can save resources and reagents by reusing standardized fragments generated by Librator. We defined a nomenclature for all fragments for easier inventory management. Each fragment name is composed of three parts: gene segment subtype (H1–H18, N1–N11), fragment number (F1–F4 for HA, F1–F3 for NA) and a unique numerical ID. For example, H3-F1-0001 denotes a gene fragment at position 1 (first fragment) generated from an H3 sequence with an ID 0001. We designed a SQL table for Librator for inventory management of all gene fragments. There are 9 keys in the fragment table: Name (prime key), Segment (HA/NA), Fragment (F1–F4), Subtype, ID, Template (template sequence name), AAseq (amino acid sequence), NTseq (nucleotide sequence), and Instock (yes/no). We also designed an interface for users to manage their fragment inventory.

### Implementation

The pipeline was primarily implemented in Python3 (version 3.7.3 for MacOS, version 3.9 for Windows 10) using PyQT5 library (version 5.13.0). The executable application was compiled from source code using Pyinstaller (version 4, https://www.pyinstaller.org/). JQuery JavaScript library (version 3.4.1, https://jquery.com/), pyecharts library (version 1.8.1, https://pyecharts.org/) and matplotlib library (version 3.1.1, https://matplotlib.org/) were used to generate figures and integrative HTML sequence viewers. Local databases were generated by sqlite3 (https://docs.python.org/3/library/sqlite3.html), a Python version of SQLite (version 3.33.0, https://www.sqlite.org/); remote databases were generated by MySQL (version 8.0, https://www.mysql.com/). The entire project was developed using PyCharm CE community (version 2019.2, https://www.jetbrains.com/pycharm/) integrated development environment. We integrated two sequence aligners MUSCLE (version 3.8.31, https://www.drive5.com/muscle/) and Clustal Omega (version 1.2.3, http://www.clustal.org/omega/) for multiple sequence alignment, H1/H3 numbering alignment, fragment alignment and mutation identification ^38, 39^. We also implemented an interface for users to visualize their sequences on 3D structures using PyMOL (version 2.3.2, https://pymol.org/) and UCSF Chimera (https://www.cgl.ucsf.edu/chimera/), to generate a maximum-likelihood tree using RAxML (version 8.0.0, https://raxml-ng.vital-it.ch/), and to visualize a phylogenetic tree using phylotree.js library (http://phylotree.hyphy.org/)^21, 22, 40^. Sequence logos of selected sequences were generated by WebLogo (version 3.7.1, https://weblogo.berkeley.edu/) ^20^. Landsacpe of multiple sequence alignment is generated by html2canvas (https://html2canvas.hertzen.com/) and Python3. The codon optimization functions is powered by DNA Chisel (version 3.2.6, https://github.com/Edinburgh-Genome-Foundry/DnaChisel). The crystal-structure-based HA numbering system is adopted from a public repository (https://github.com/bloomlab/HA_numbering) with some modifications.

### Software and code availability

Librator is freely hosted online (https://wilsonimmunologylab.github.io/Librator/). Tutorials are available from a Wilson Lab GitHub page (https://wilsonimmunologylab.github.io/Librator/), a pdf format user guide is also available for downloading. The source code is also available from GitHub (https://github.com/WilsonImmunologyLab/Librator).

We provide executable version of this software for two dominated operating systems: Windows 10 and MacOS. The MacOS version of this software is compiled under macOS Mojave (version 10.14.6) and has been tested under macOS Mojave (version 10.14.6), macOS Catalina (version 10.15.2) and macOS Big Sur (version 11.2.3). The Windows 10 version of this software is compiled under Windows 10 Home (OS build 19042.867) and has been tested under the same system.

This Python-based software is also transferable and can be compiled under other systems (e.g. ubuntu) from source code.

## Acknowledgements

We would like to thank Dr. Jesse Bloom for his assistance and suggestions for this project.

## Author Contributions

L.L. designed the model, implemented the software, performed computational analyses, and wrote the manuscript. O.S., J.J.G., S.C. and Y.F. tested software, improved software design, and revised the manuscript. H.L.D. and C.T.S. tested software and improved software design. N.Z. and M.H. performed experimental validations. P.C.W. initiated and supervised the work, designed the model, implemented the software, and wrote the manuscript.

## SUPPLEMENTAL FIGURES

**Figure S1.**
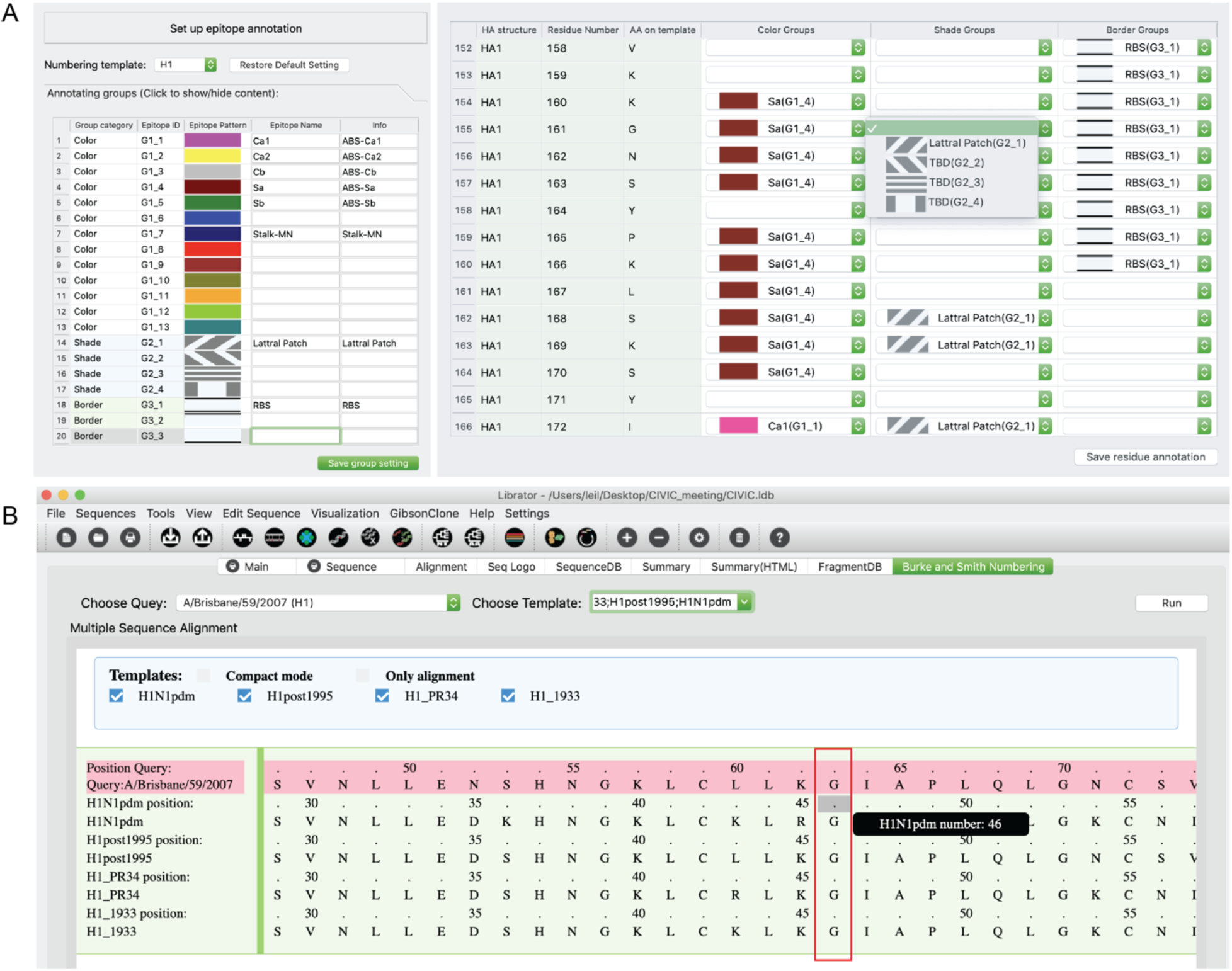
Built-in functions of Librator for influenza sequence analysis. (**A**) User customizable epitopes definition in Librator. Librator allows users to use 13 distinct colors, 4 shade patterns and 3 border styles to annotate individual residues on the sequence viewer according to their specific research interest and focus. The left panel showed GUI of defining epitopes in Librator, and the right panel showed GUI of annotating individual residues using user-defined epitopes. (**B**) Burke and Smith HA numbering scheme viewer in Librator. Users are allowed to align query sequence against multiple templates to access residue numbers on different templates.

**Figure S2.**
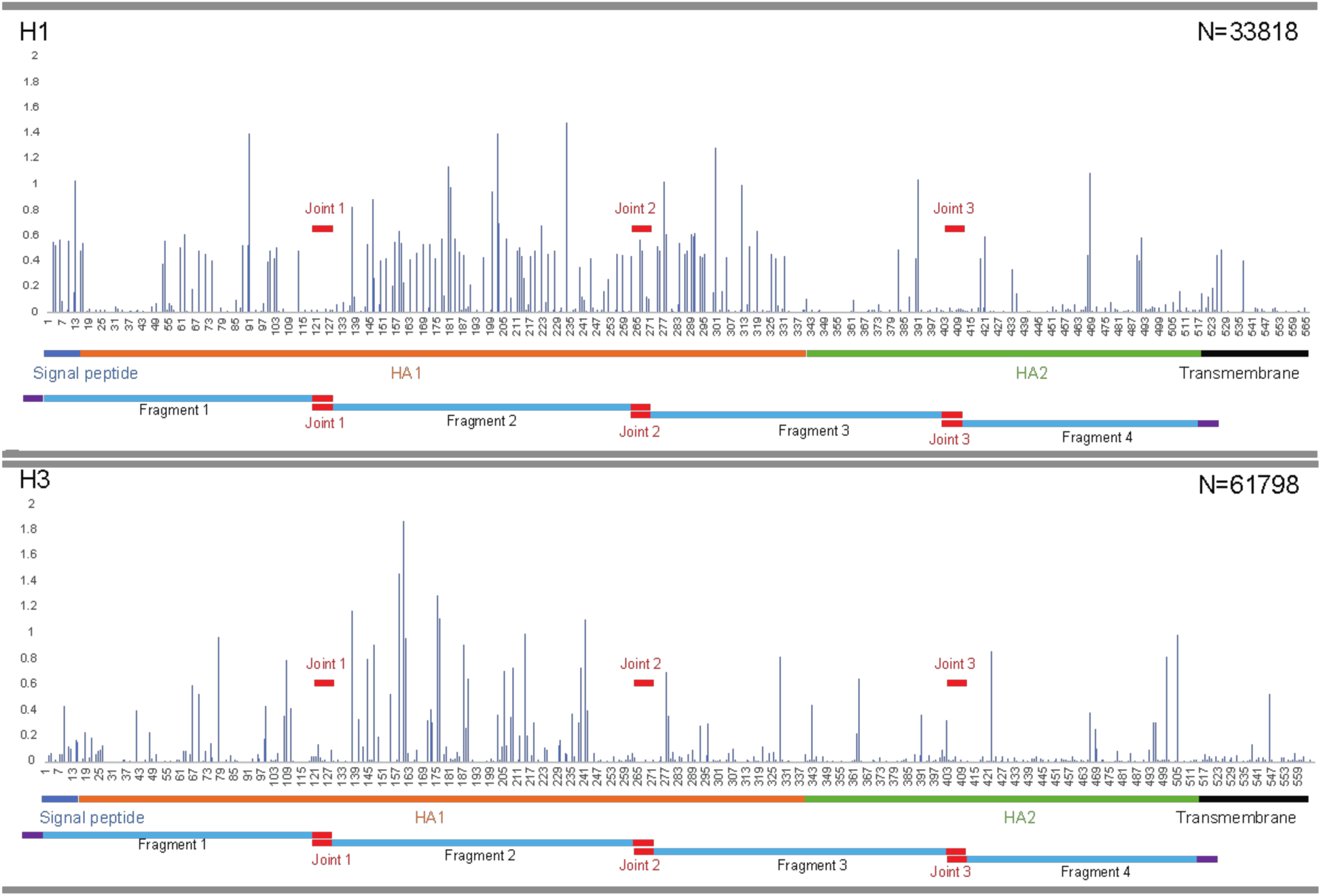
Fragment design for influenza HA proteins. HA proteins are clustered into two groups: group 1 and group 2. In Librator, all group 1 sequences are aligned to an H1 template, and all group 2 sequences are aligned to an H3 template.

**Figure S3.**
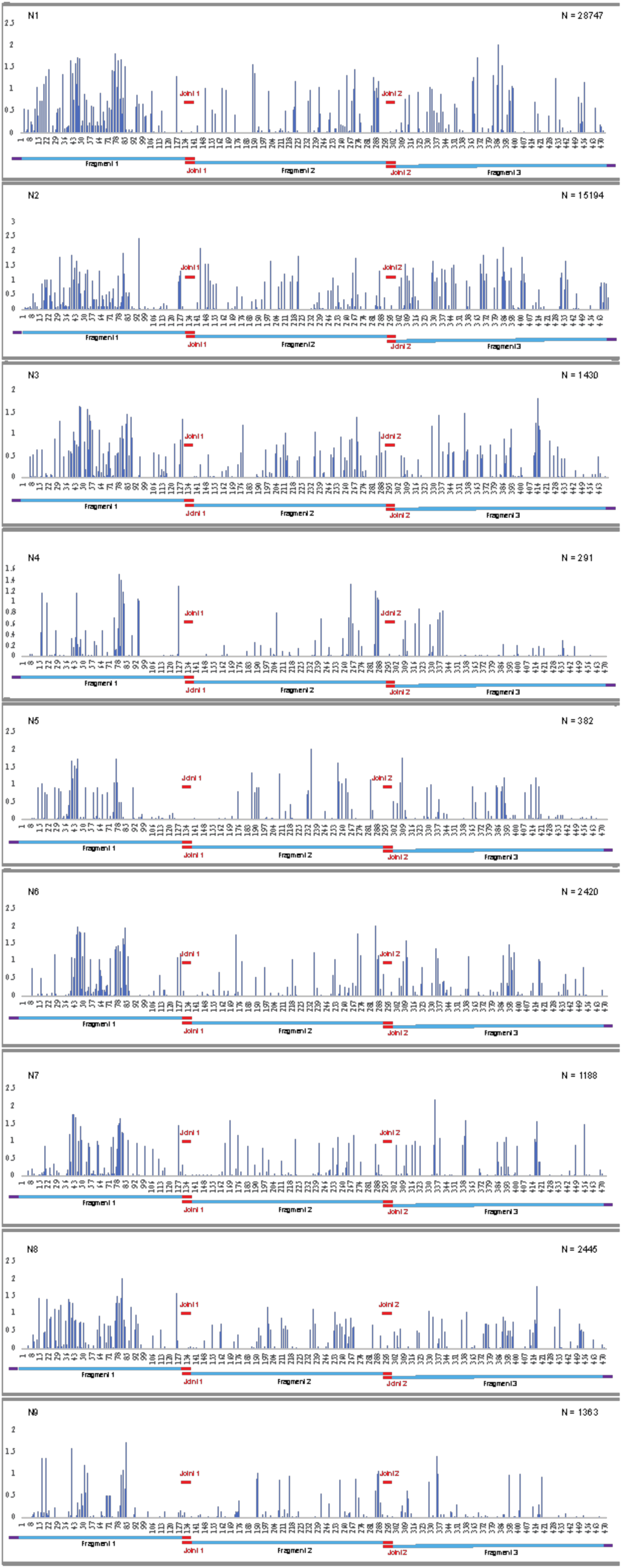
Fragment design for influenza NA proteins.

**Figure S4.**
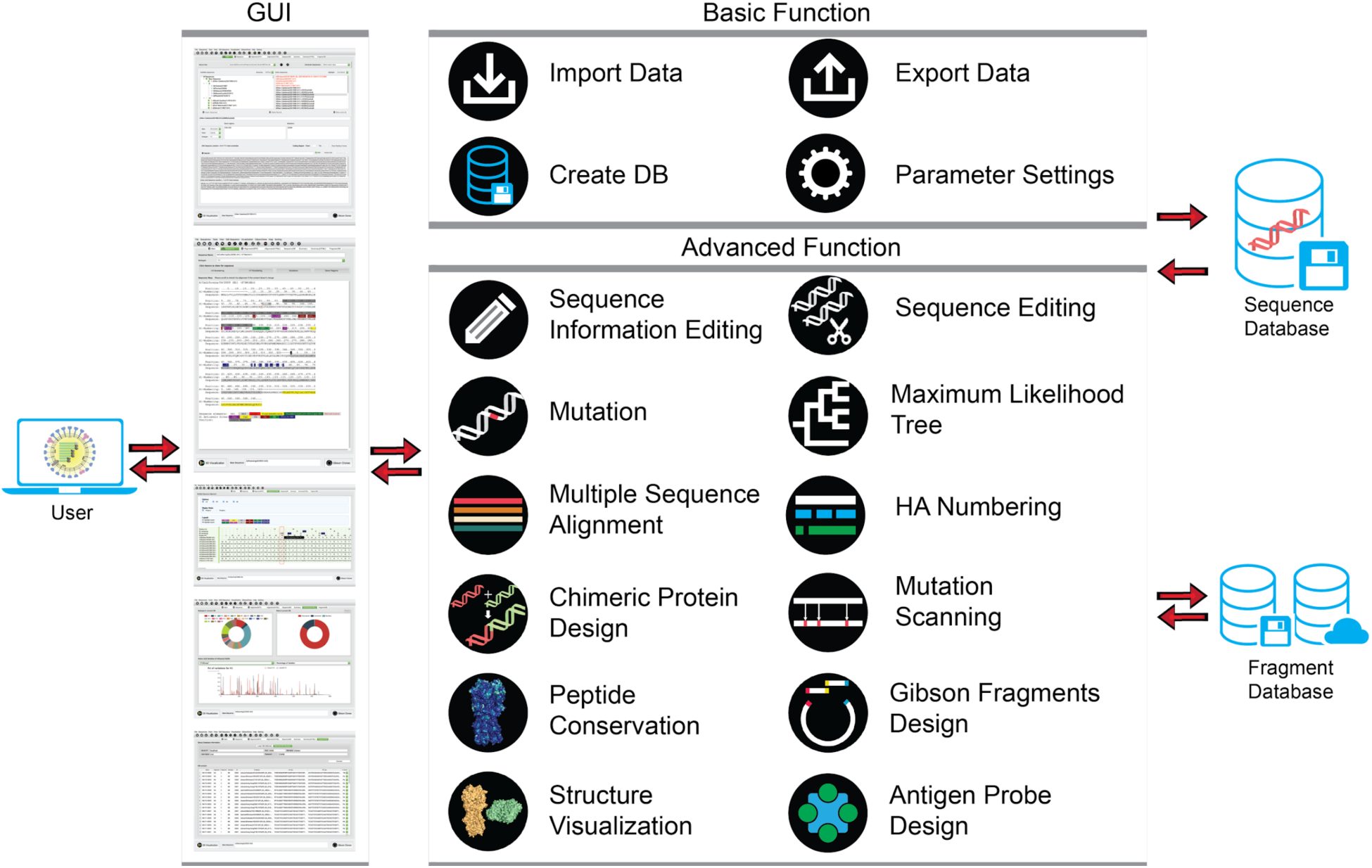
System structure and functions of Librator. Librator is comprised of a UI layer (GUI), logical layer (all functions) and data layer (SQL databases). Users are allowed to finish all operations using the GUI. All functions can be divided into two broad categories: basic function and advanced function. Basic function includes I/O operations and database (DB) operations, and advanced function includes sequence design/editing, fragment design, phylogenetic analysis and structure visualization.

**Figure S5.**
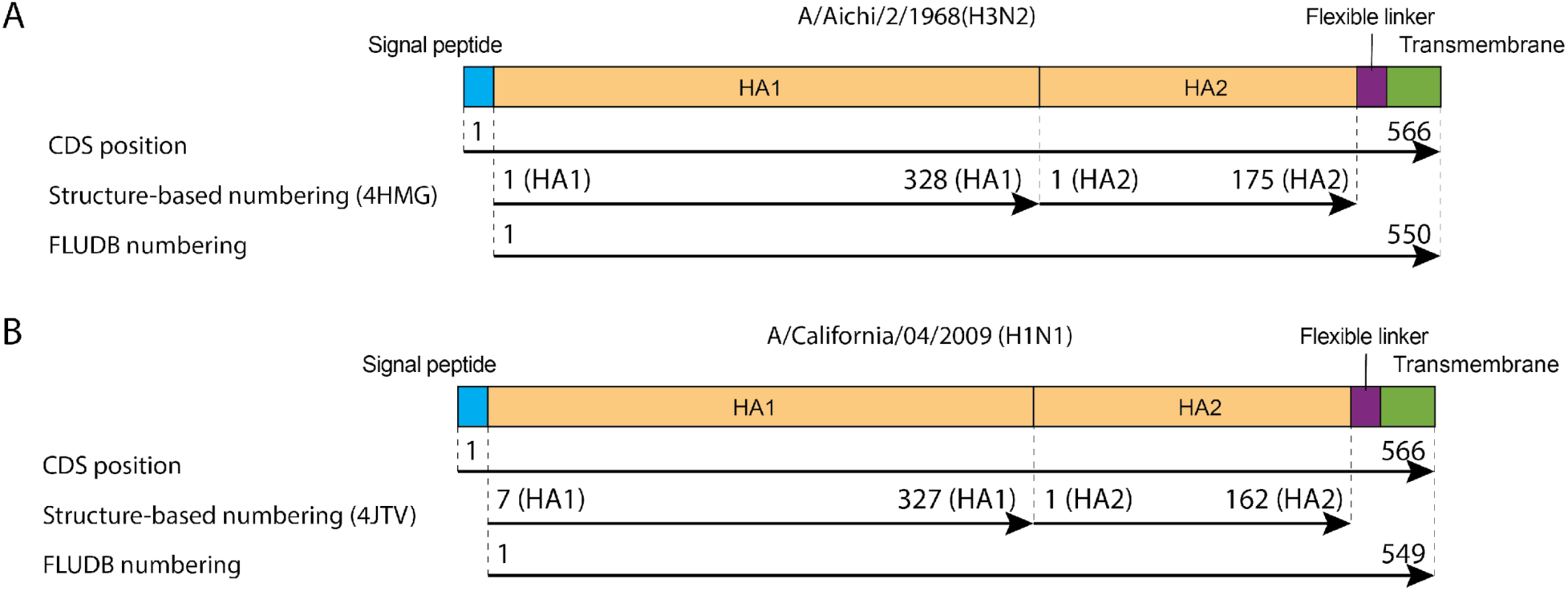
Comparison of three different HA numbering systems using a classic H1 (A/California/04/2009, H1N1) and a classic H3 (A/Aichi/2/1968, H3N2) sequence.

## SUPPLEMENTAL TABLES

**Table S1.**
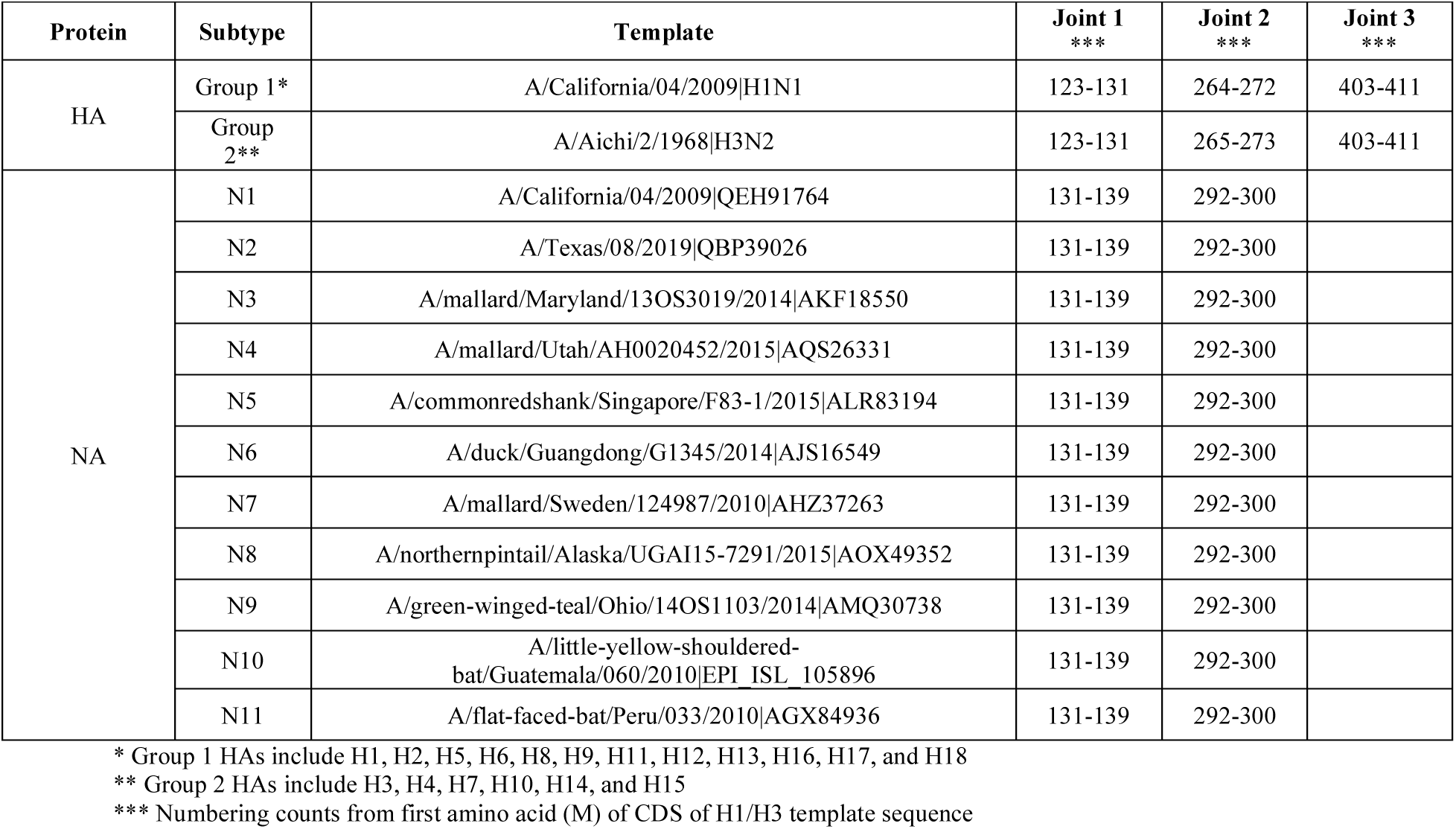
Template sequences and joint region design of HA and NA sequences in Librator.

**Table S2.**
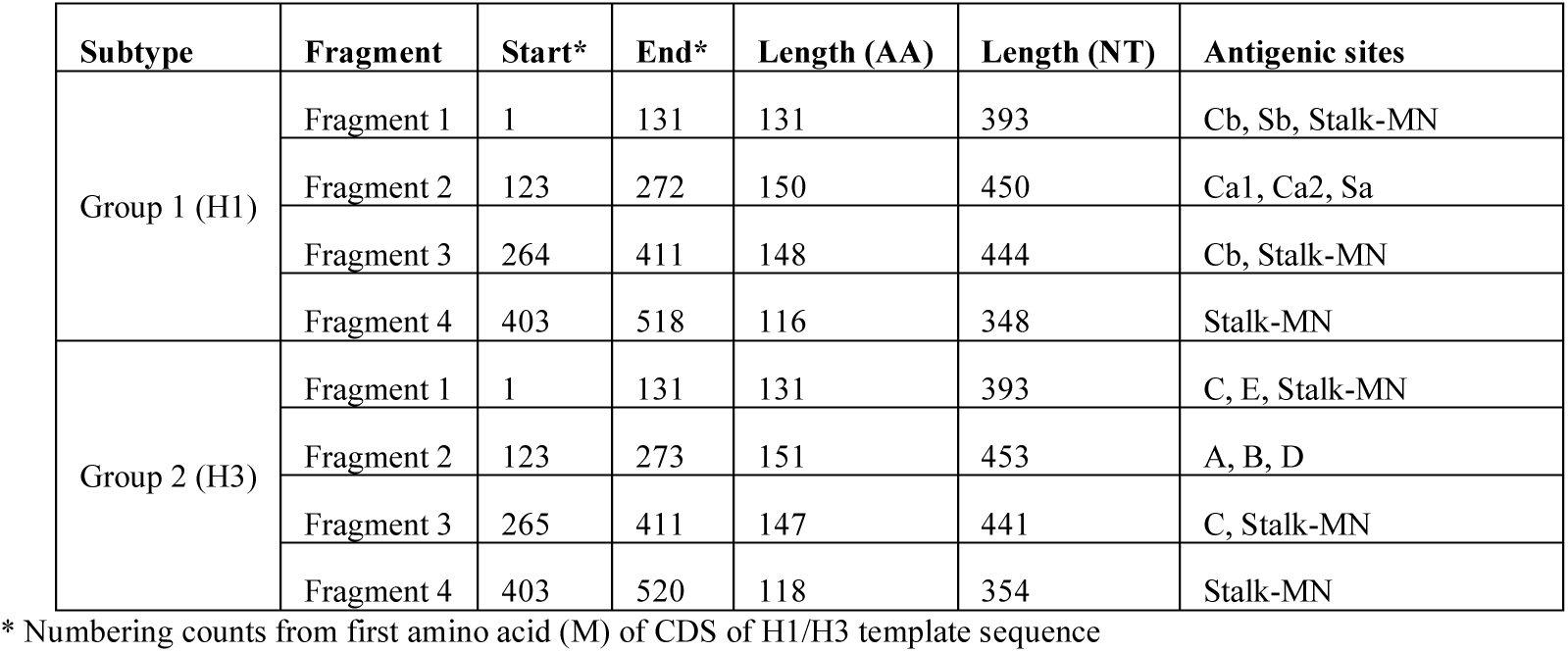
Fragment design for Group 1 HA and Group 2 HA protein sequences.

**Table S3.**
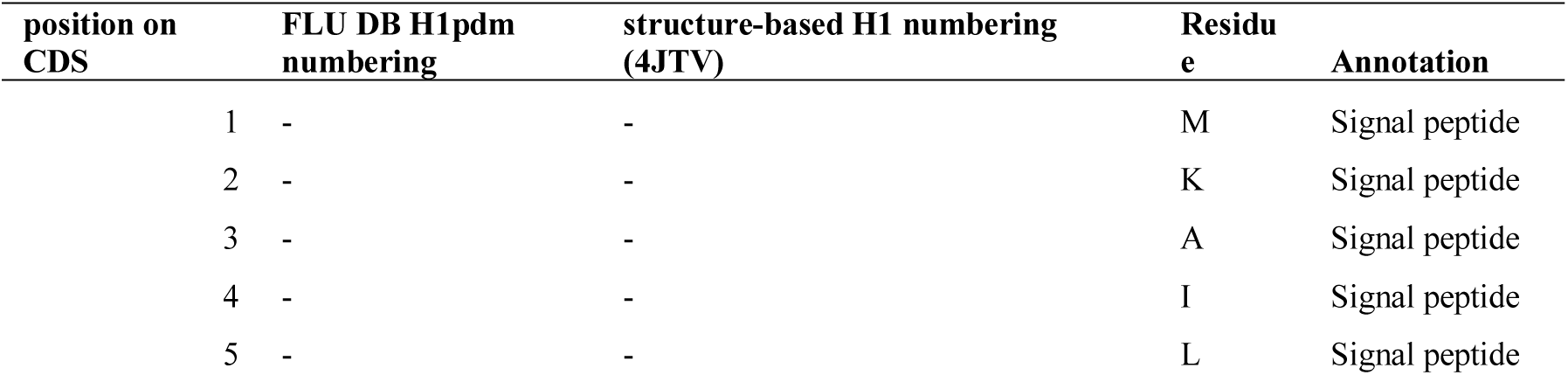
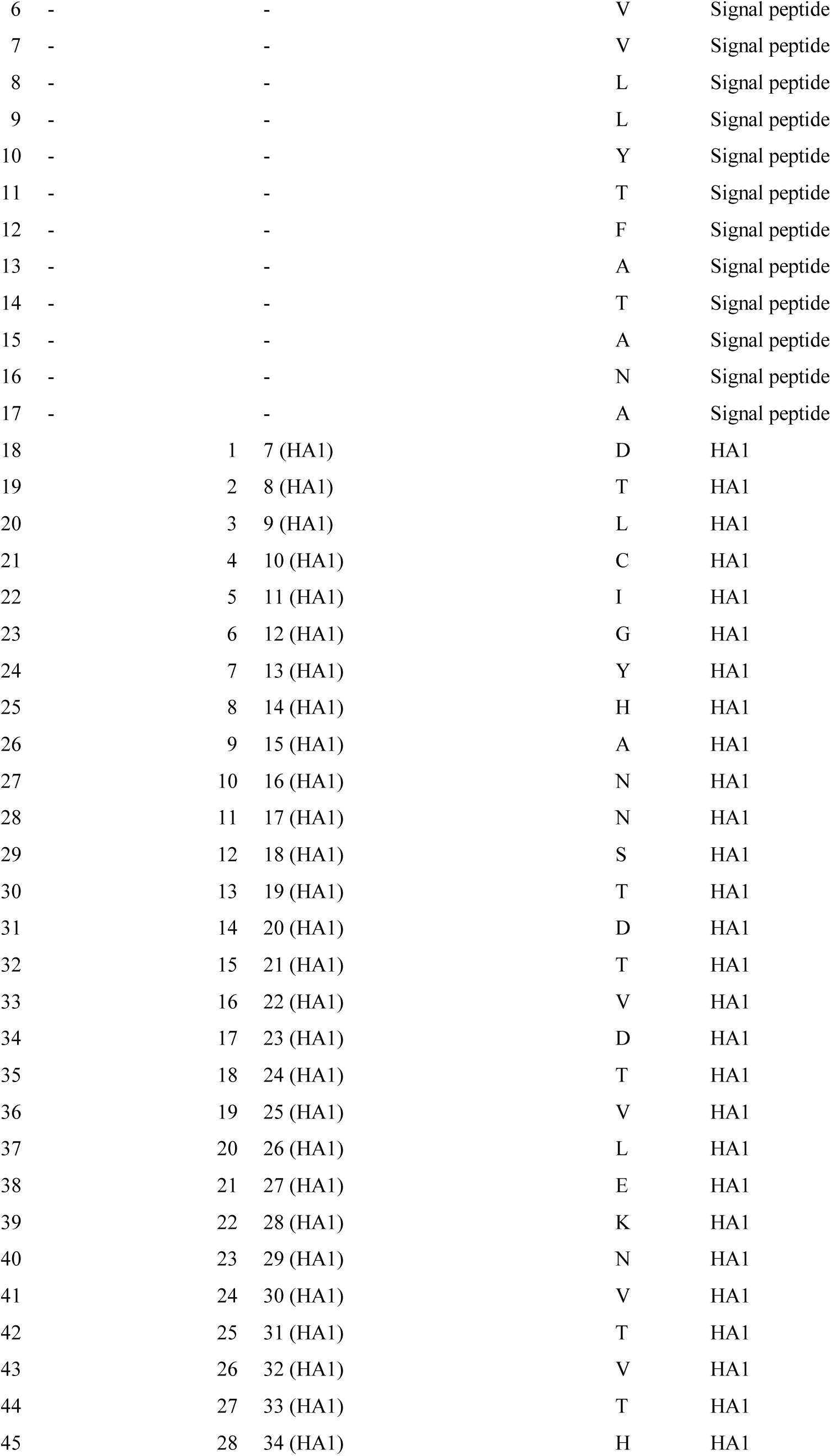
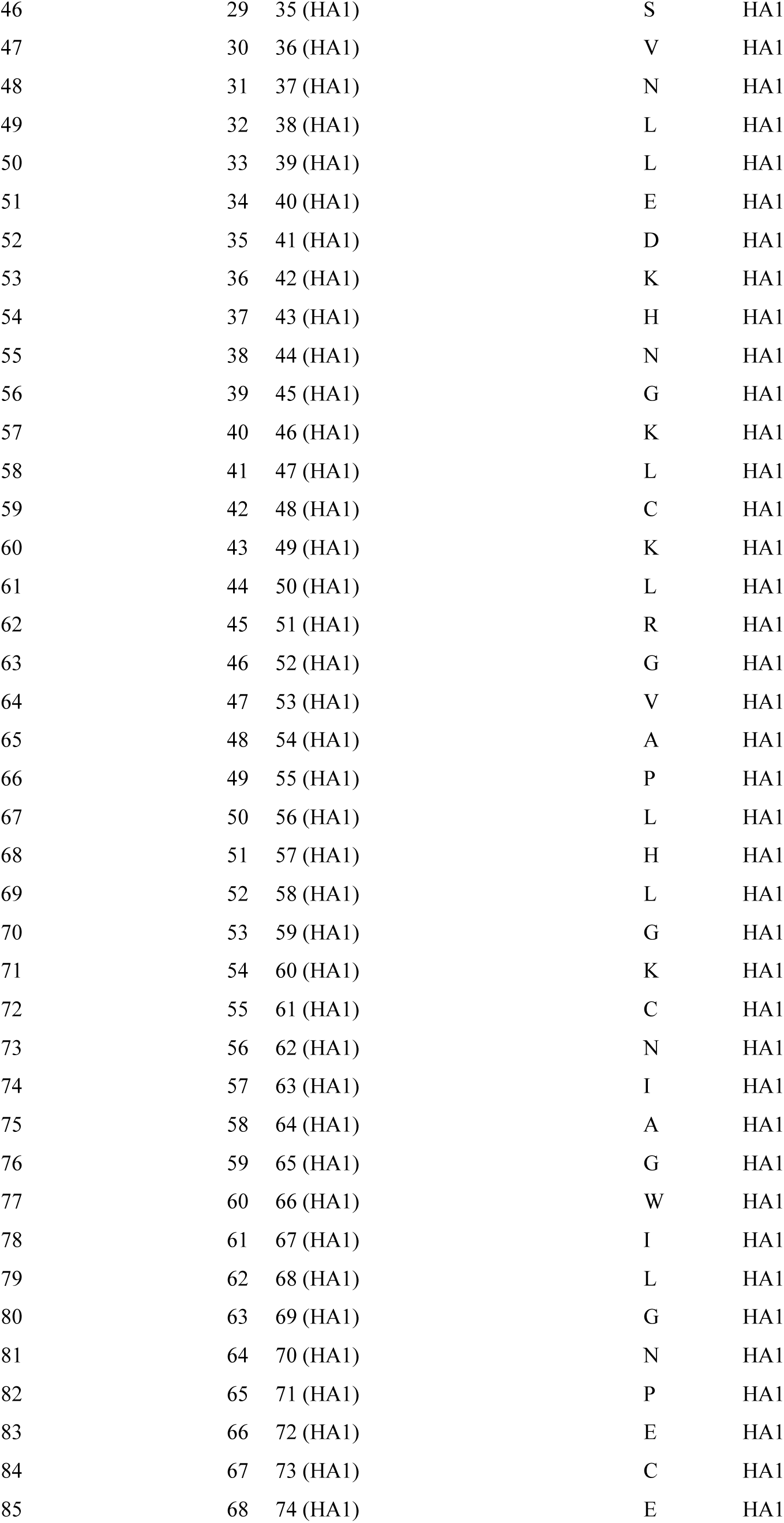
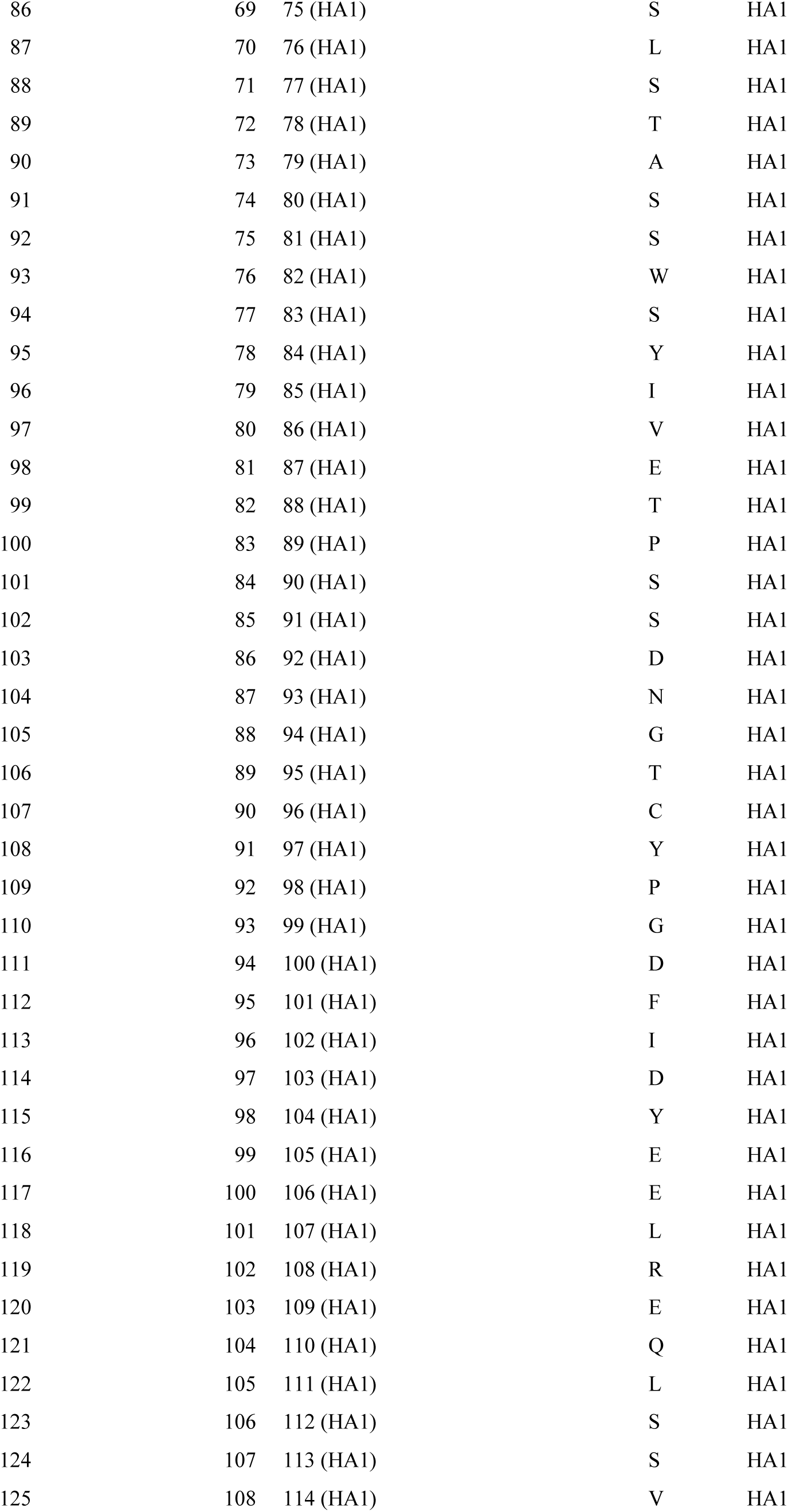
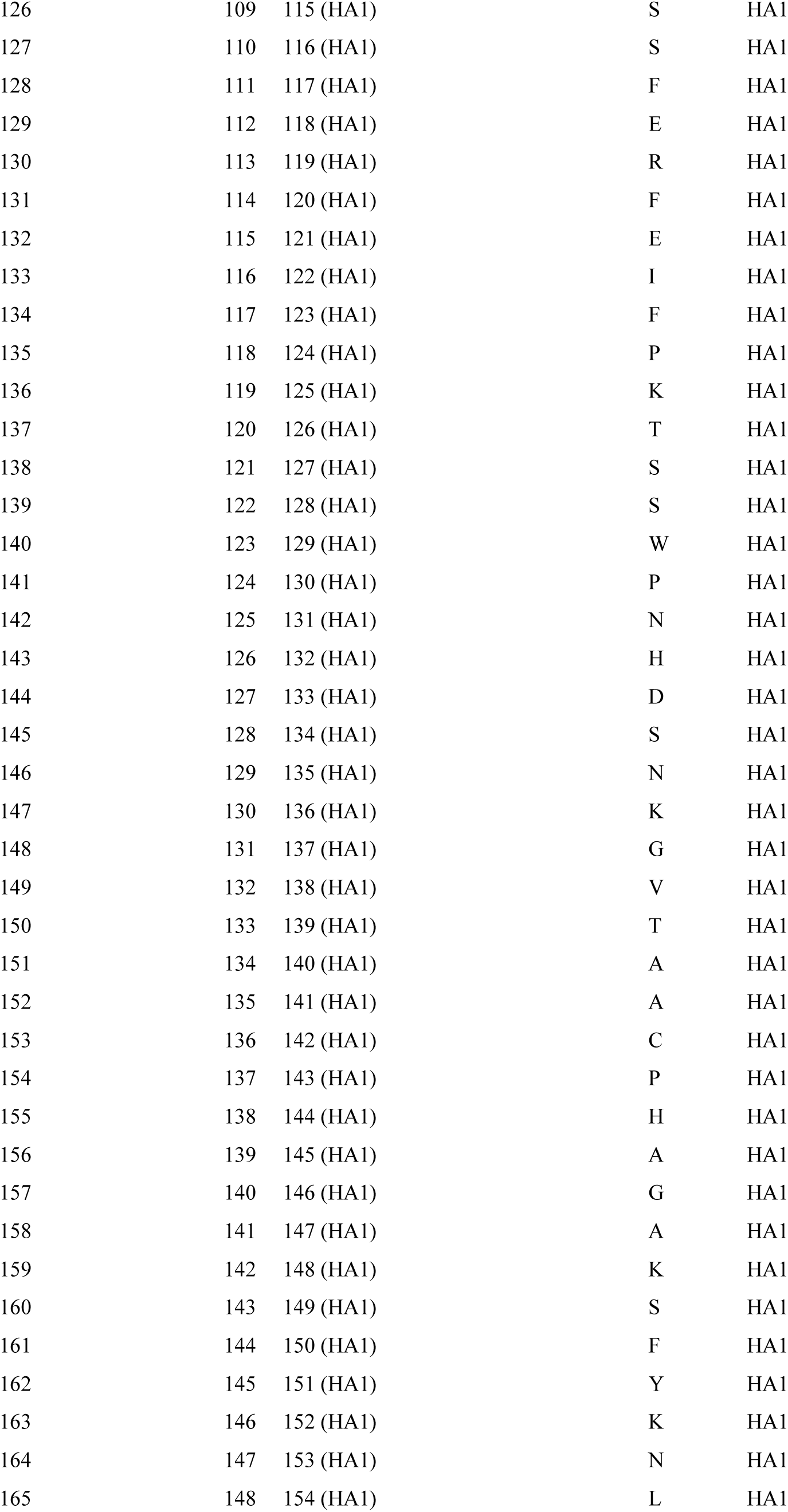
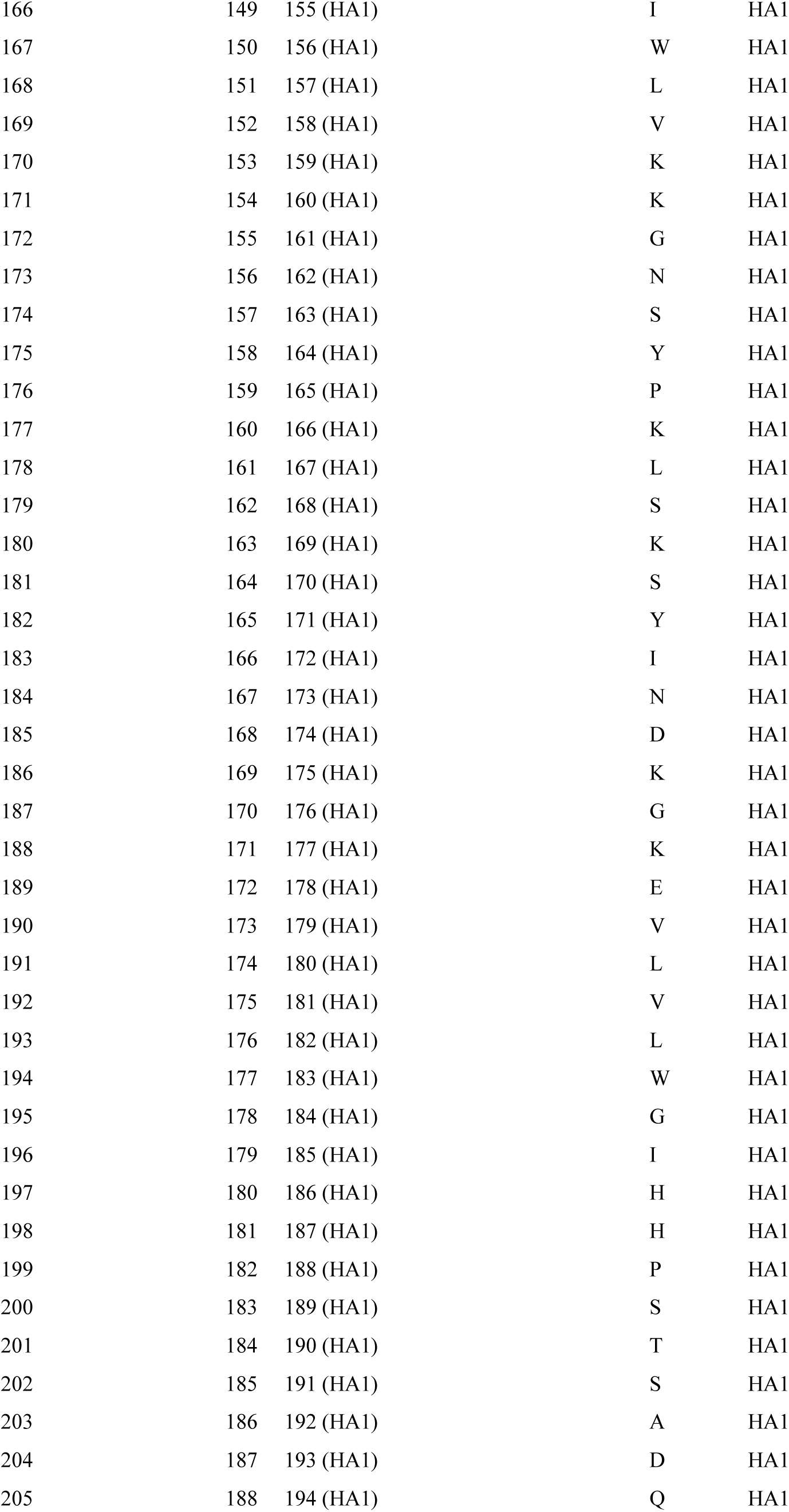
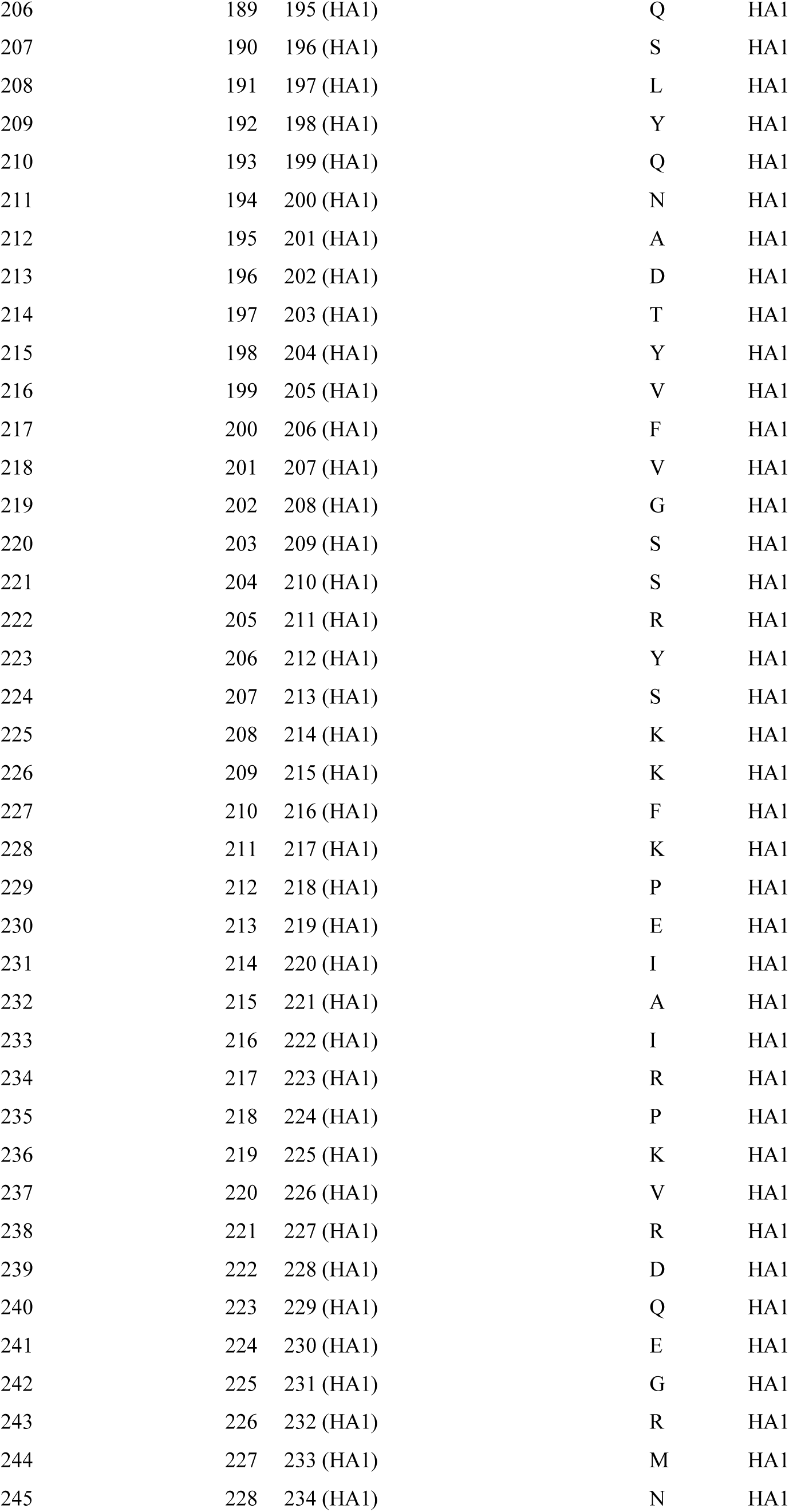
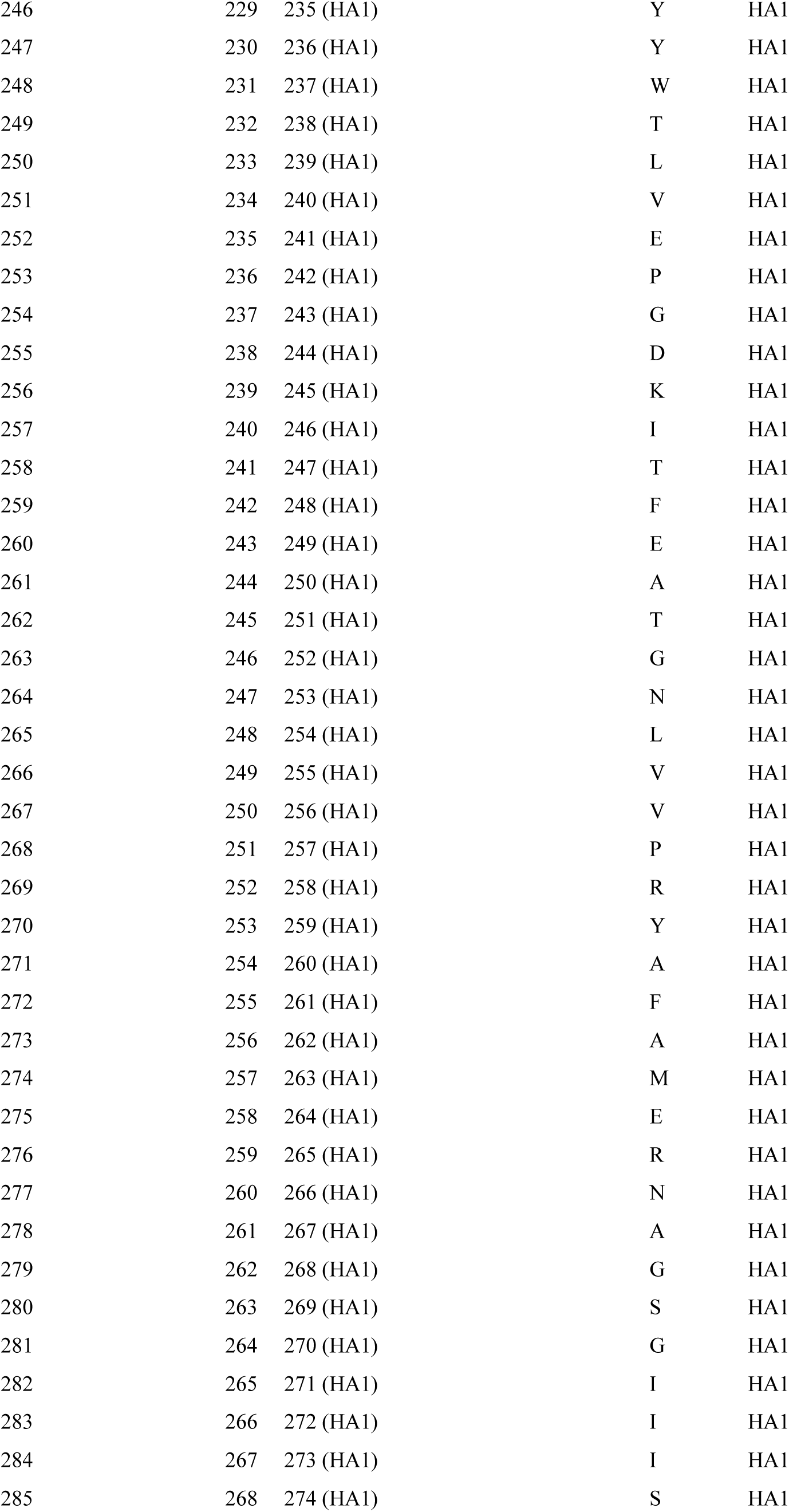
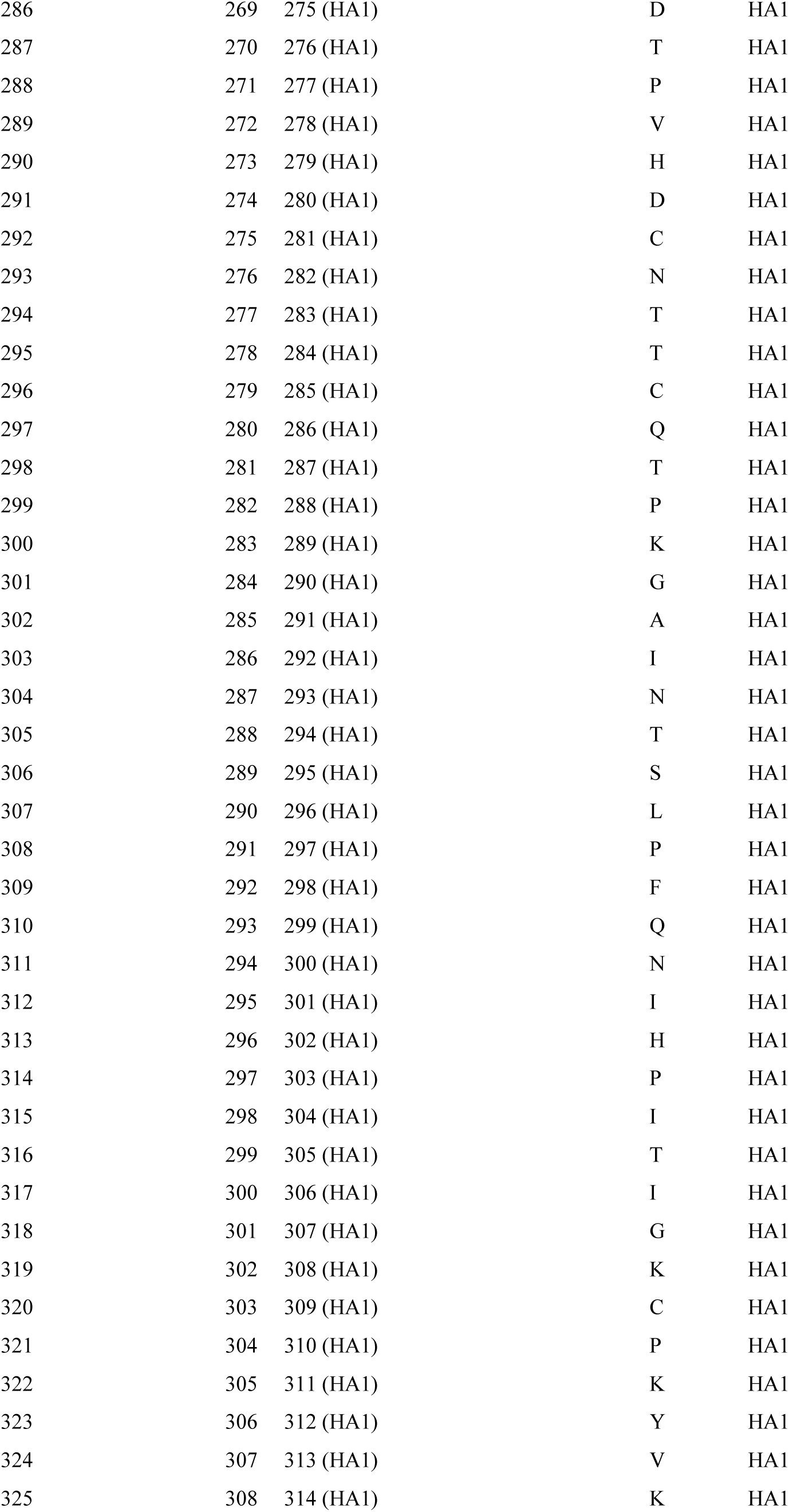
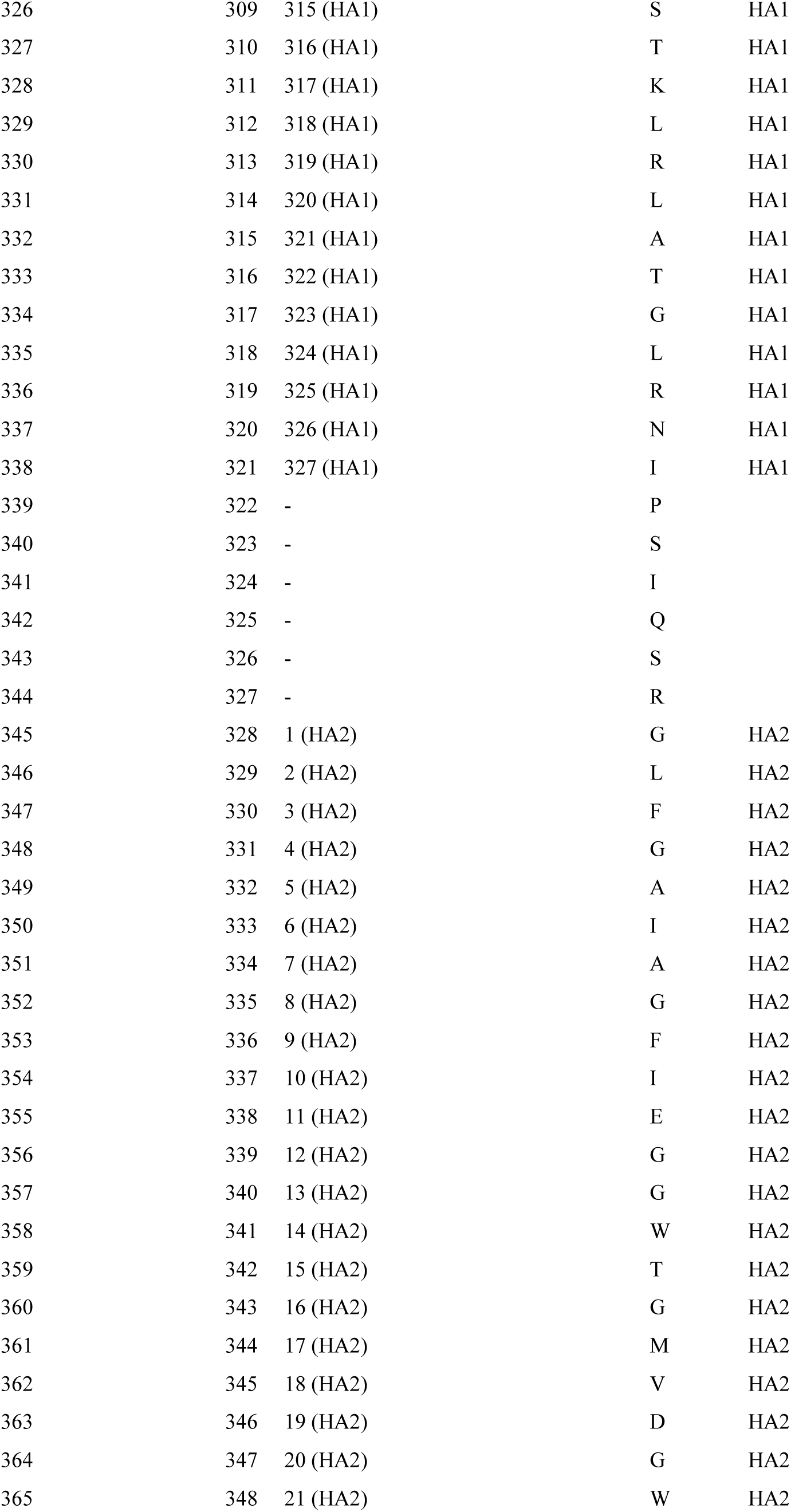
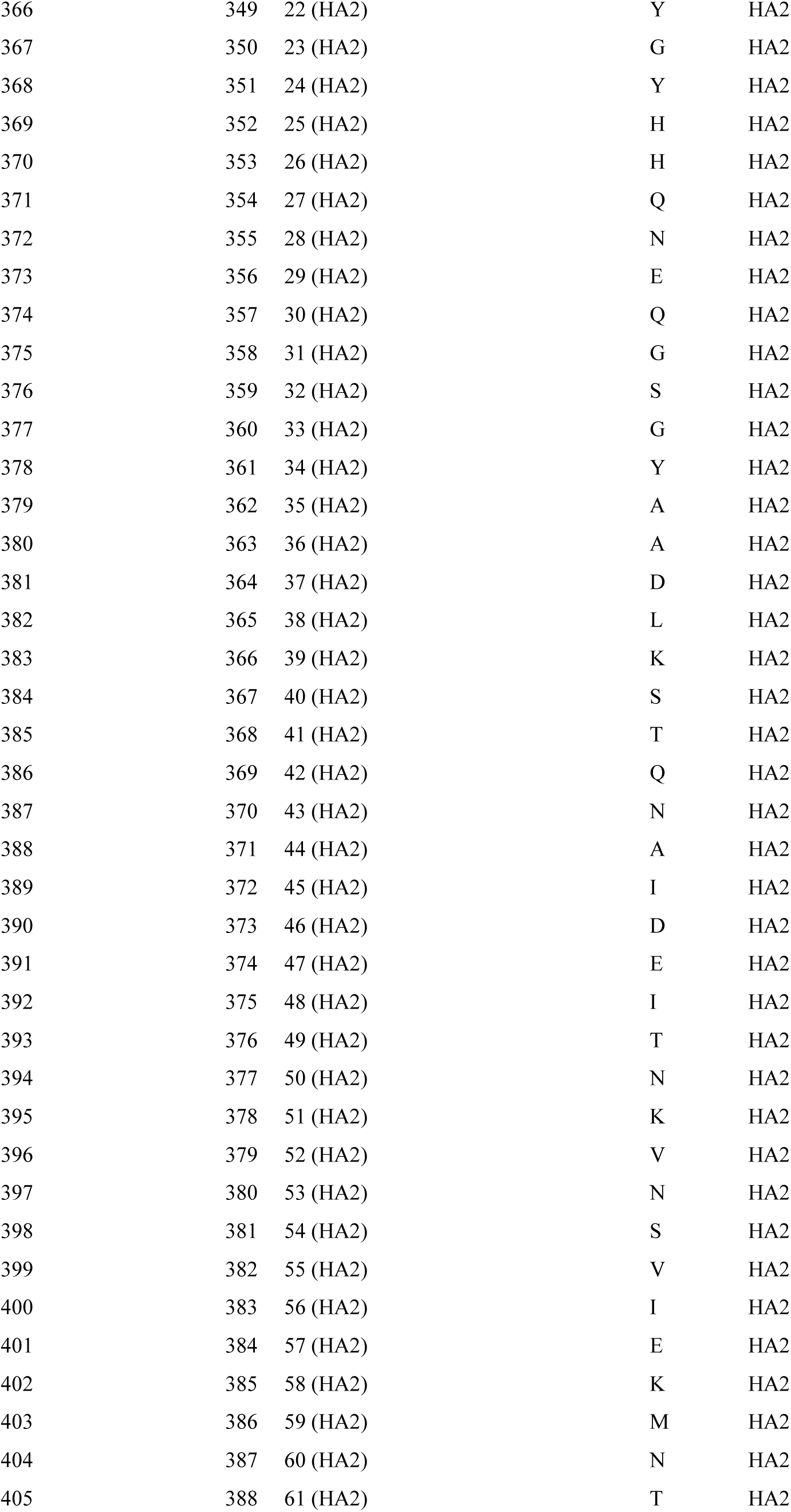
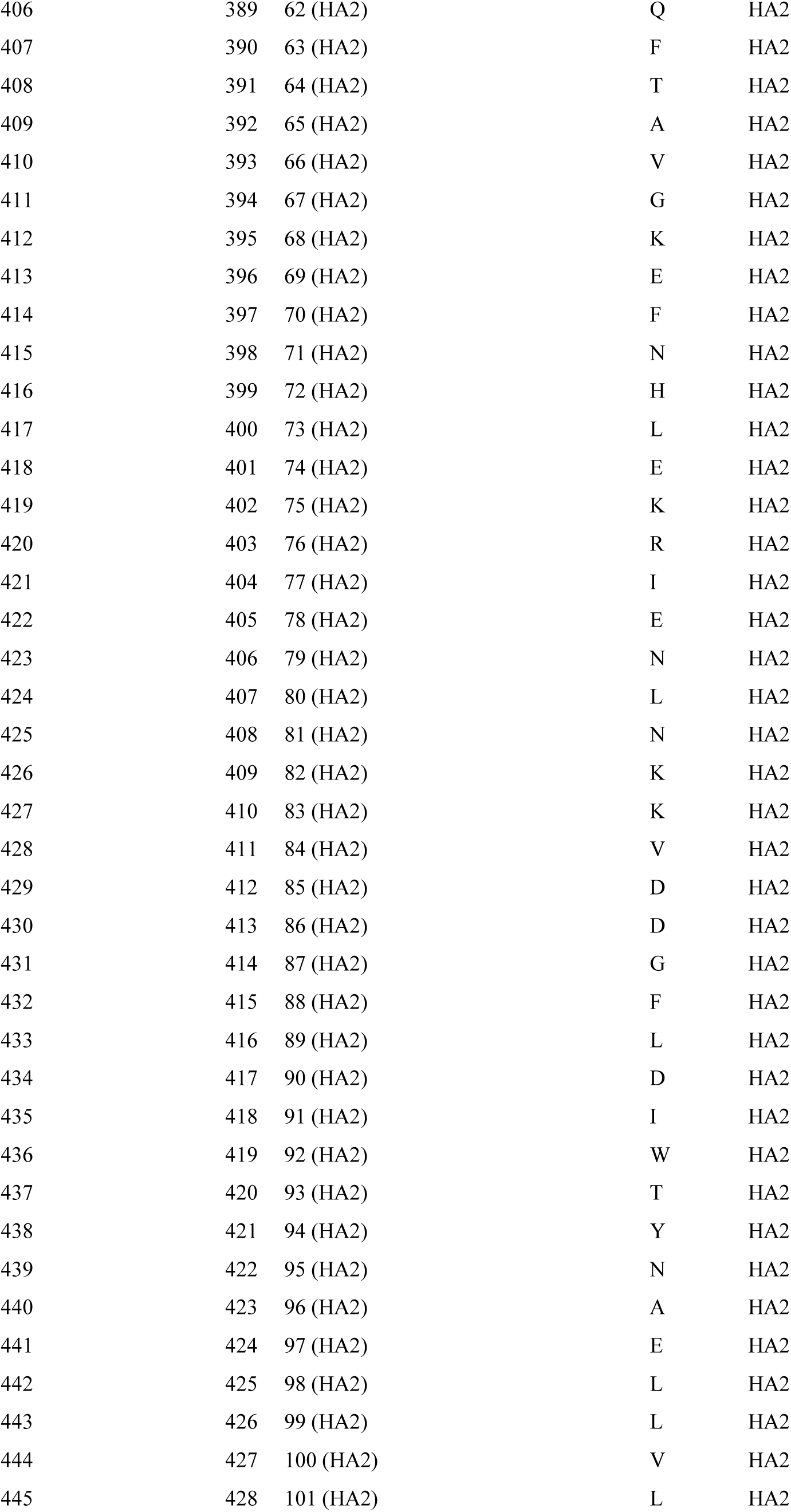
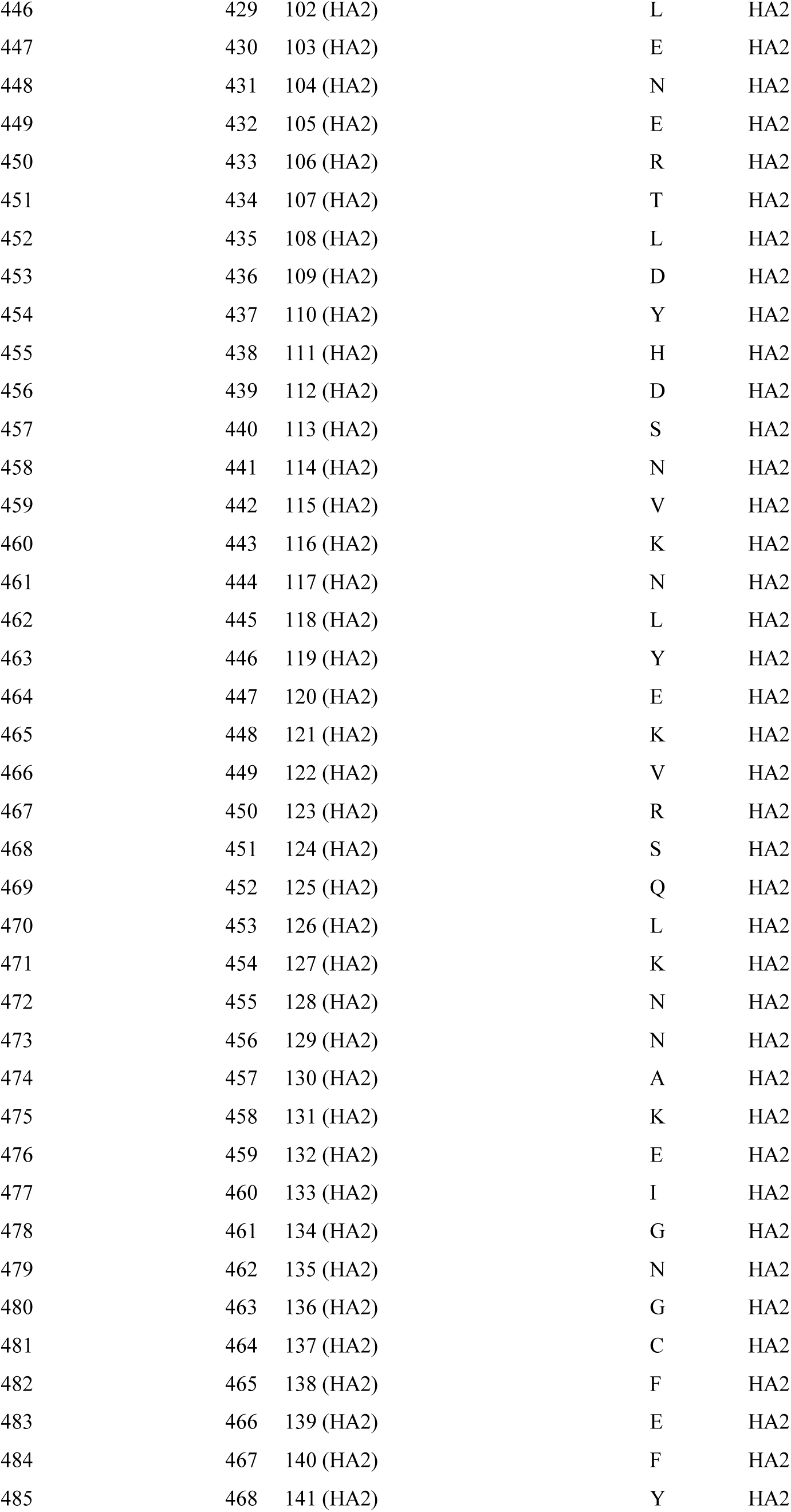
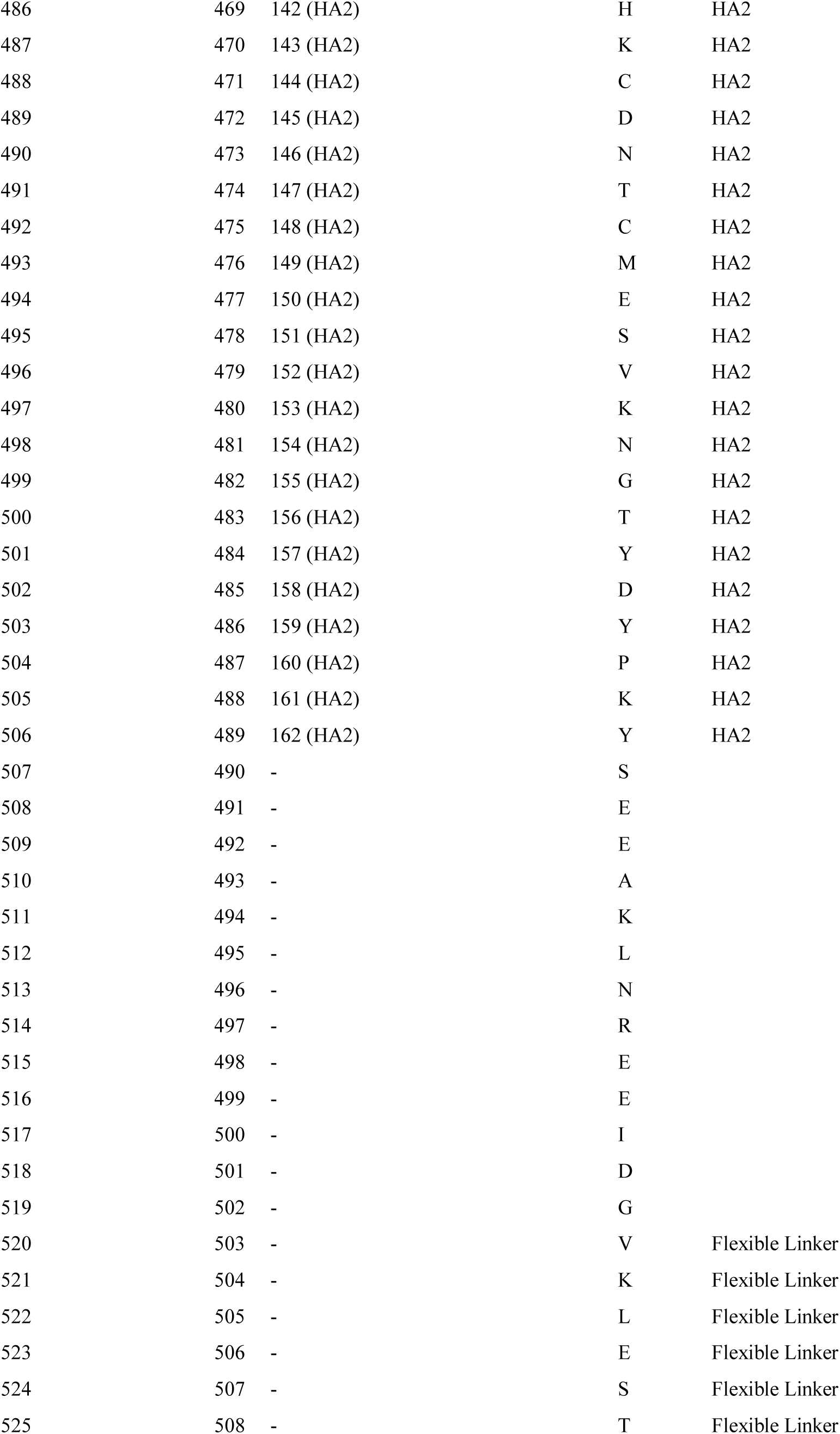
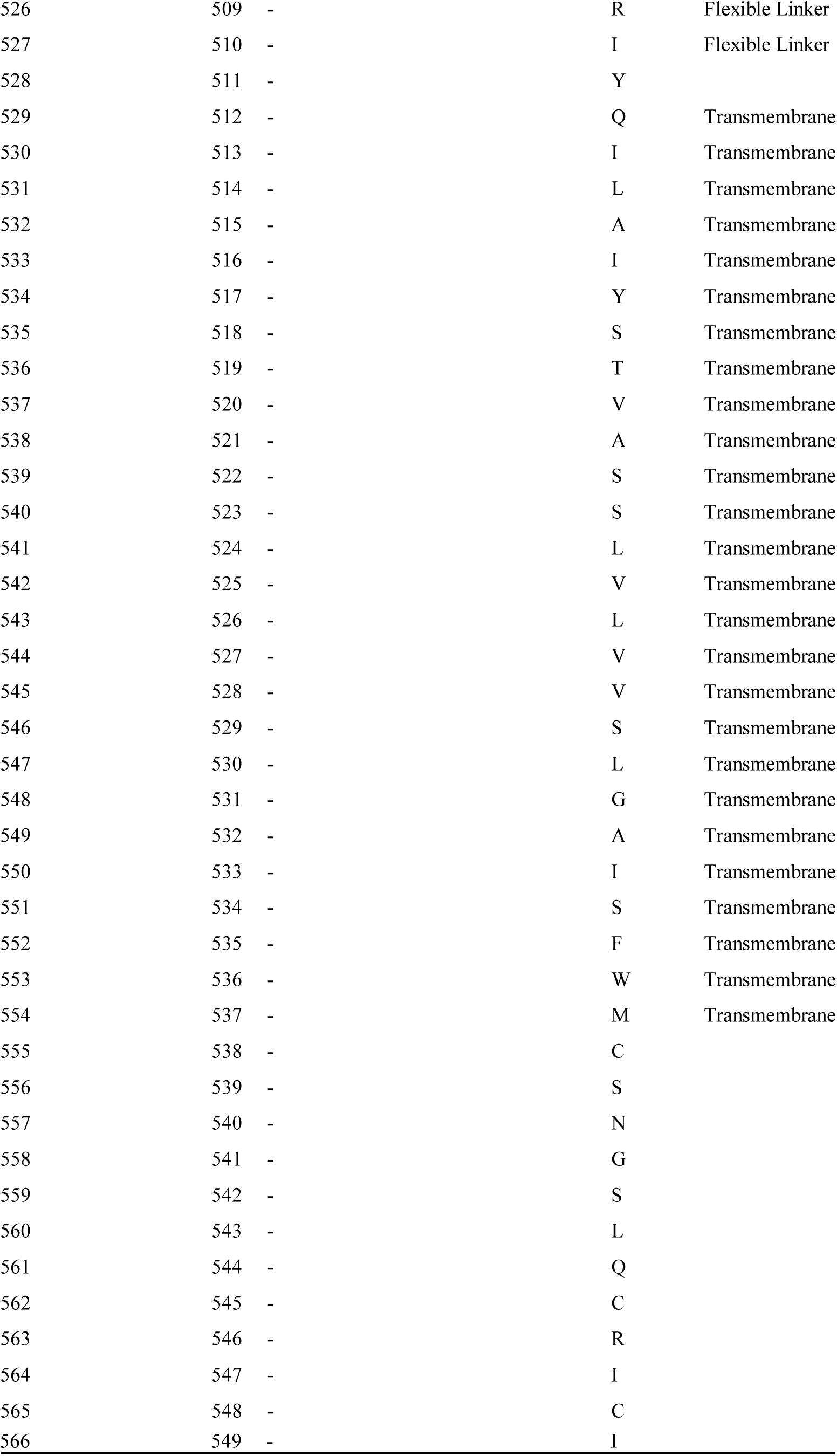
Comparison of multiple HA numbering schemes using pdm09 H1 template (A/California/04/2009, H1N1). Template for FLU DB numbering is H1N1pdm. Flexible linker and transmembrane domain were located by aligning to A/duck/Alberta/35/76(H1N1), PDB ID: 6HJR.

**Table S4.**
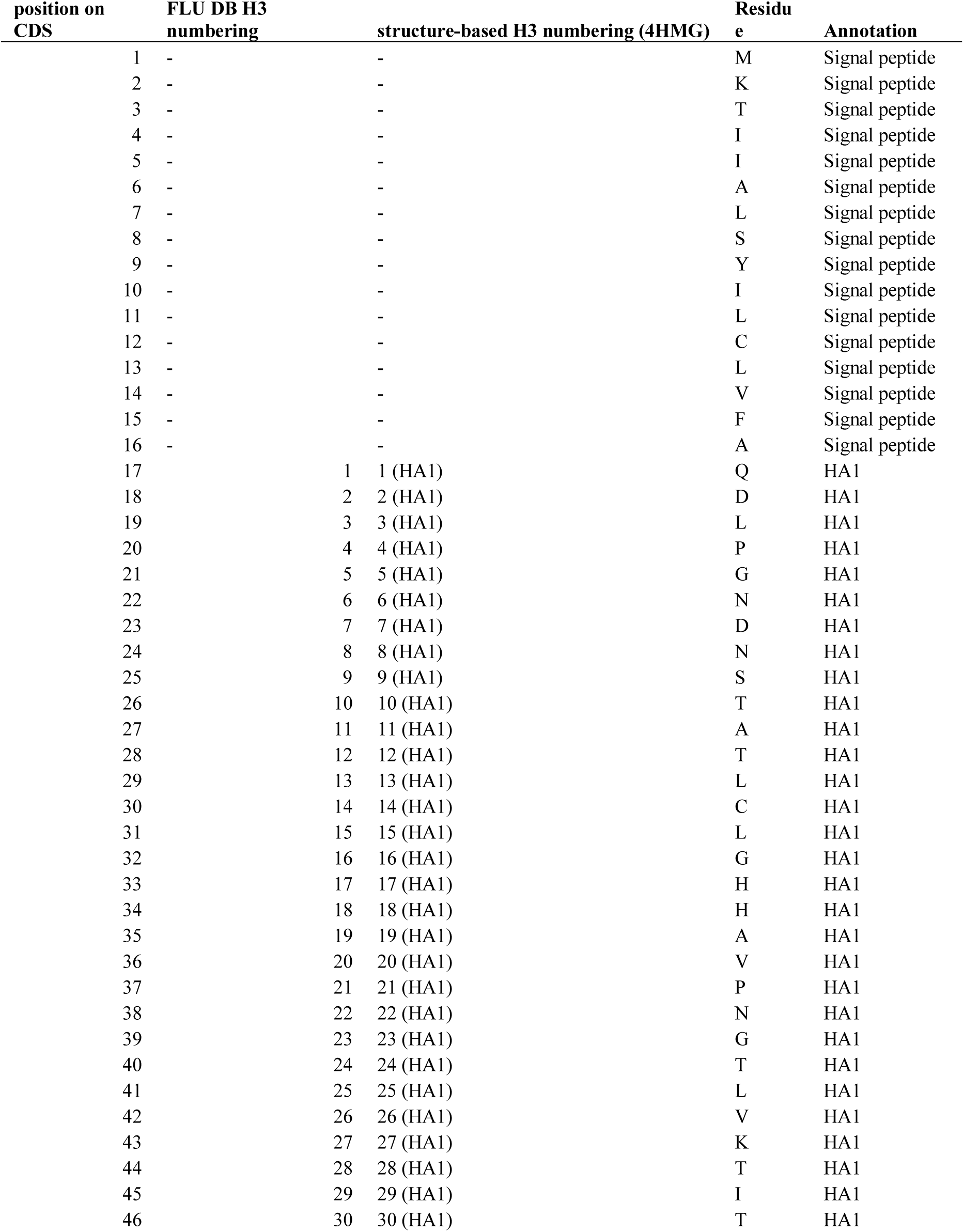
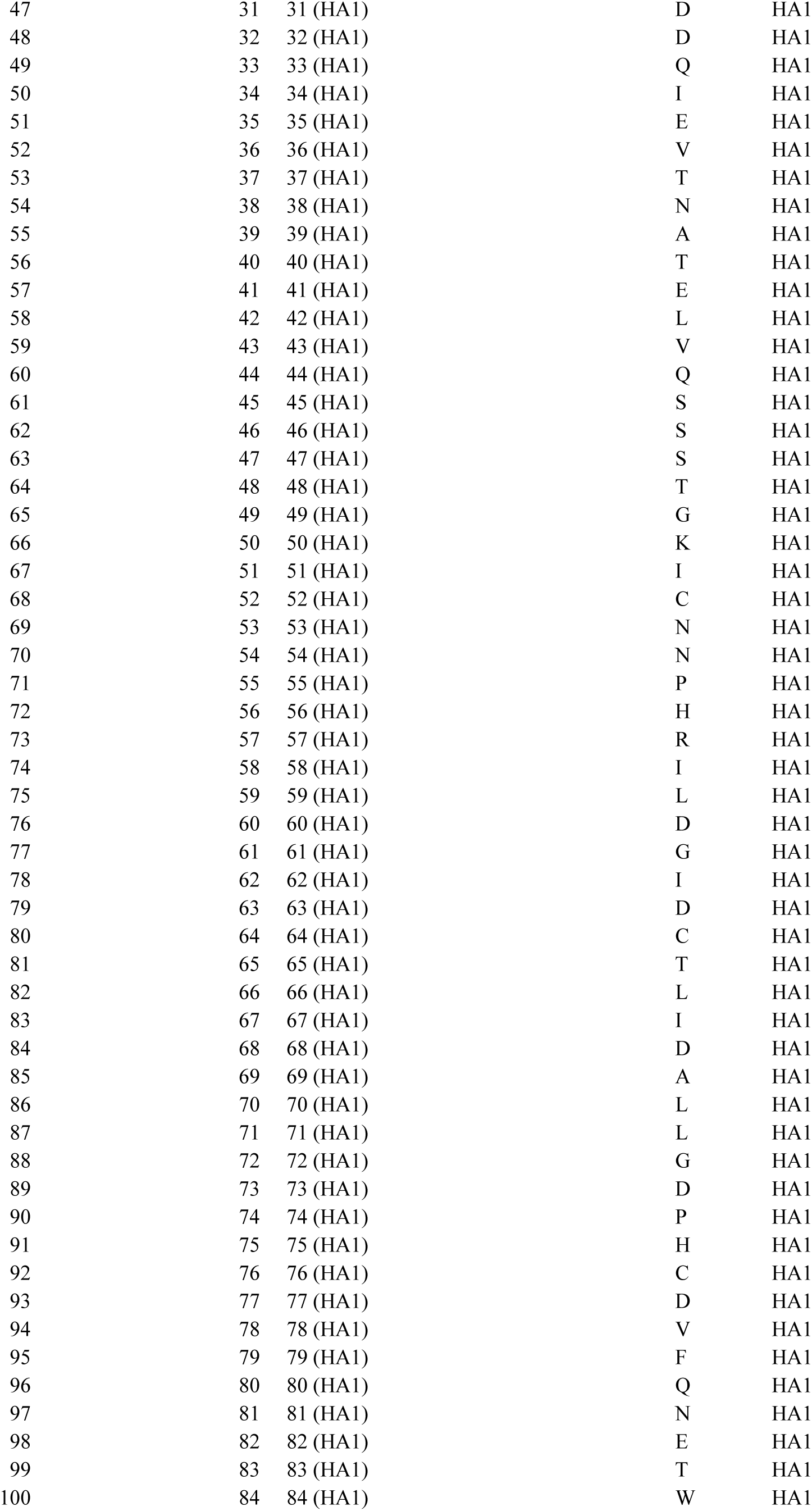
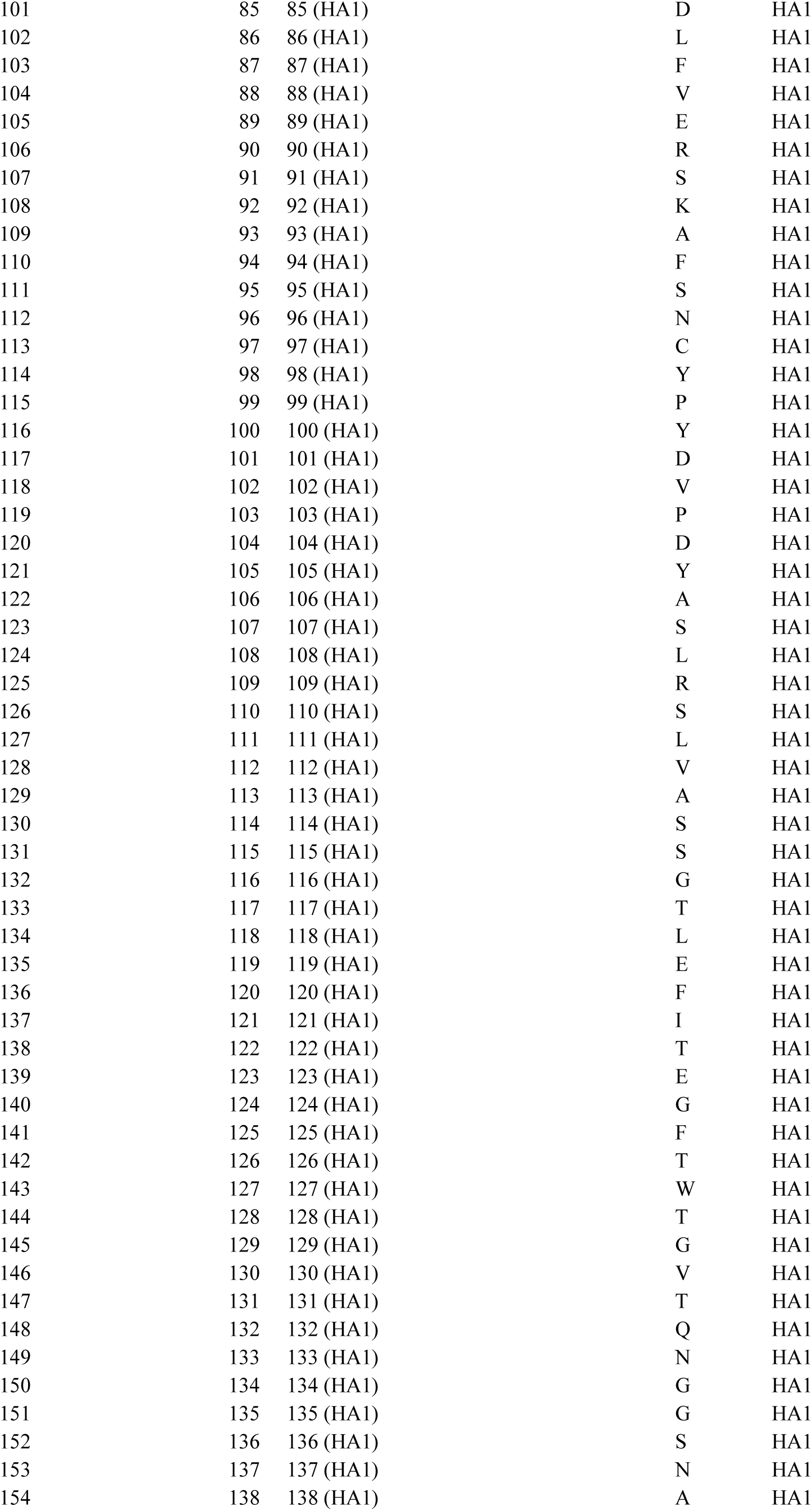
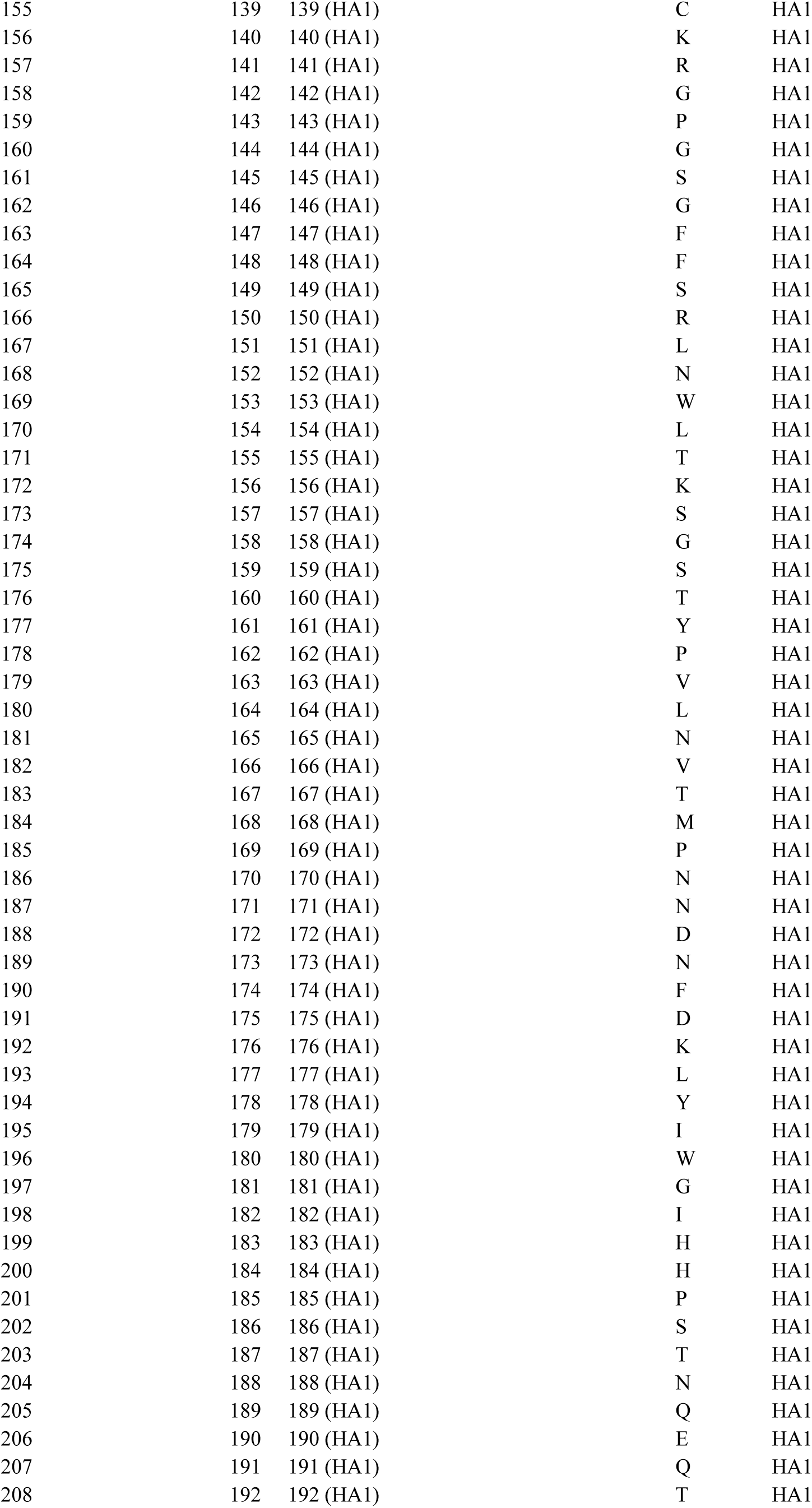
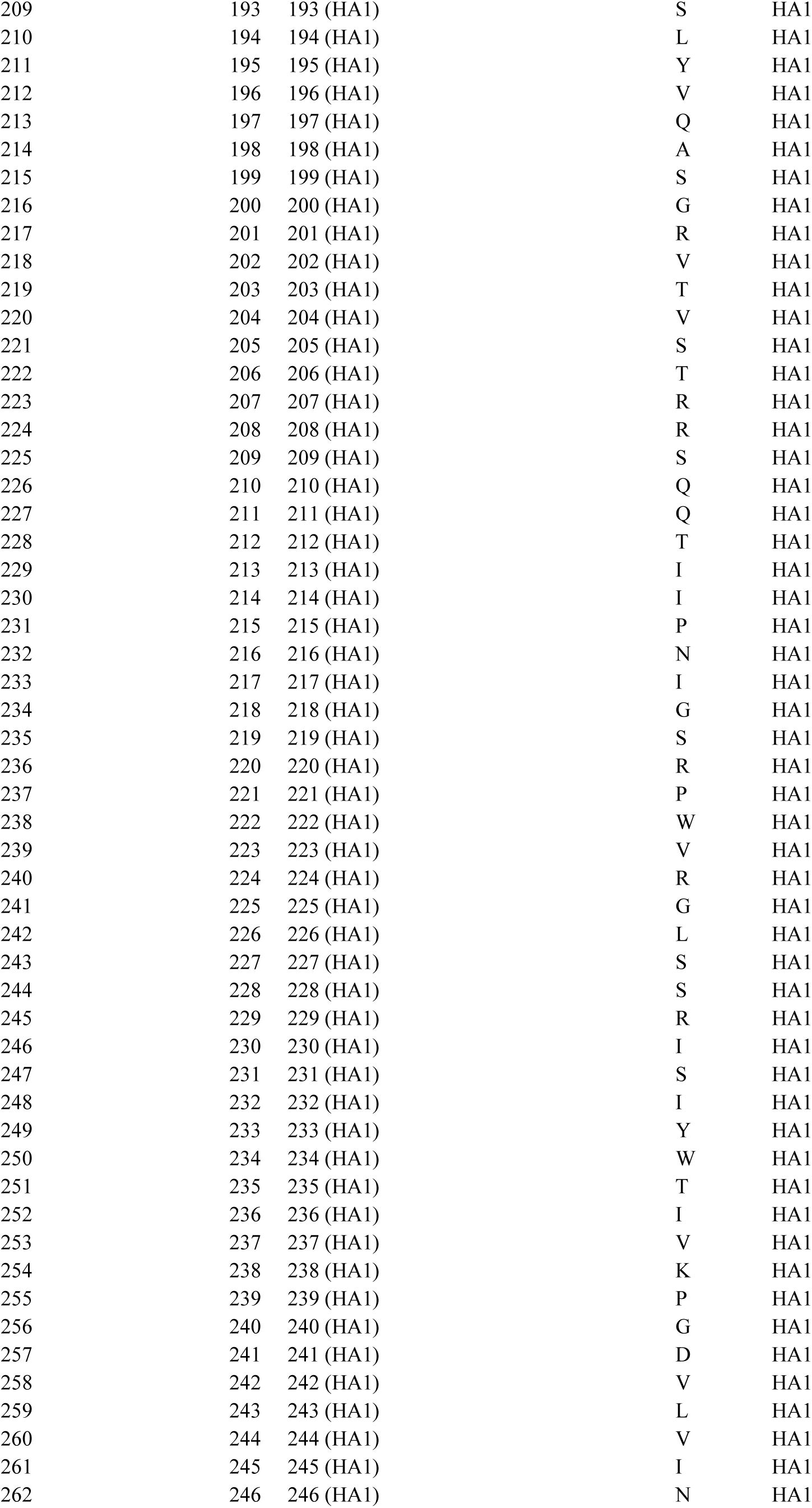
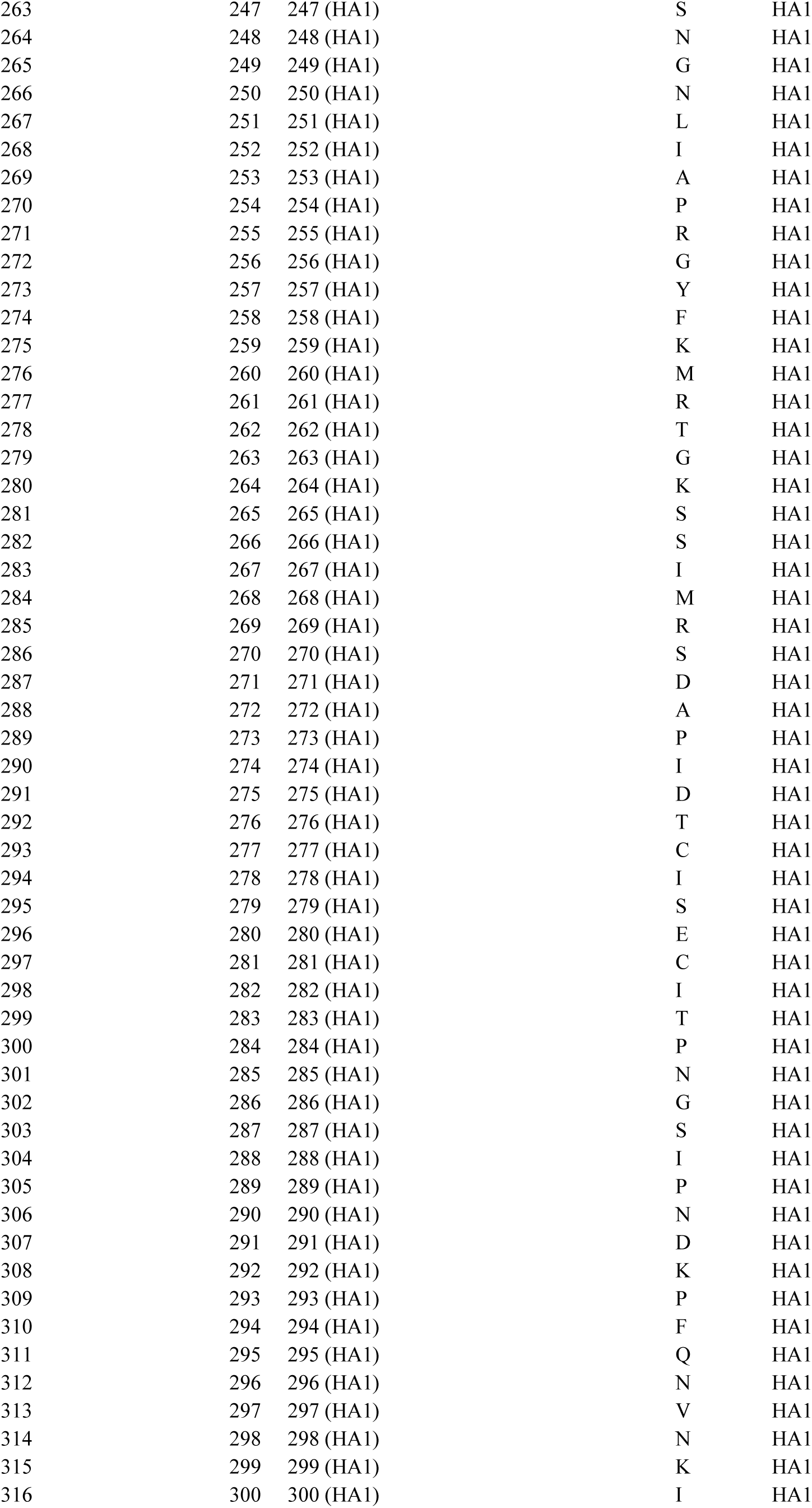
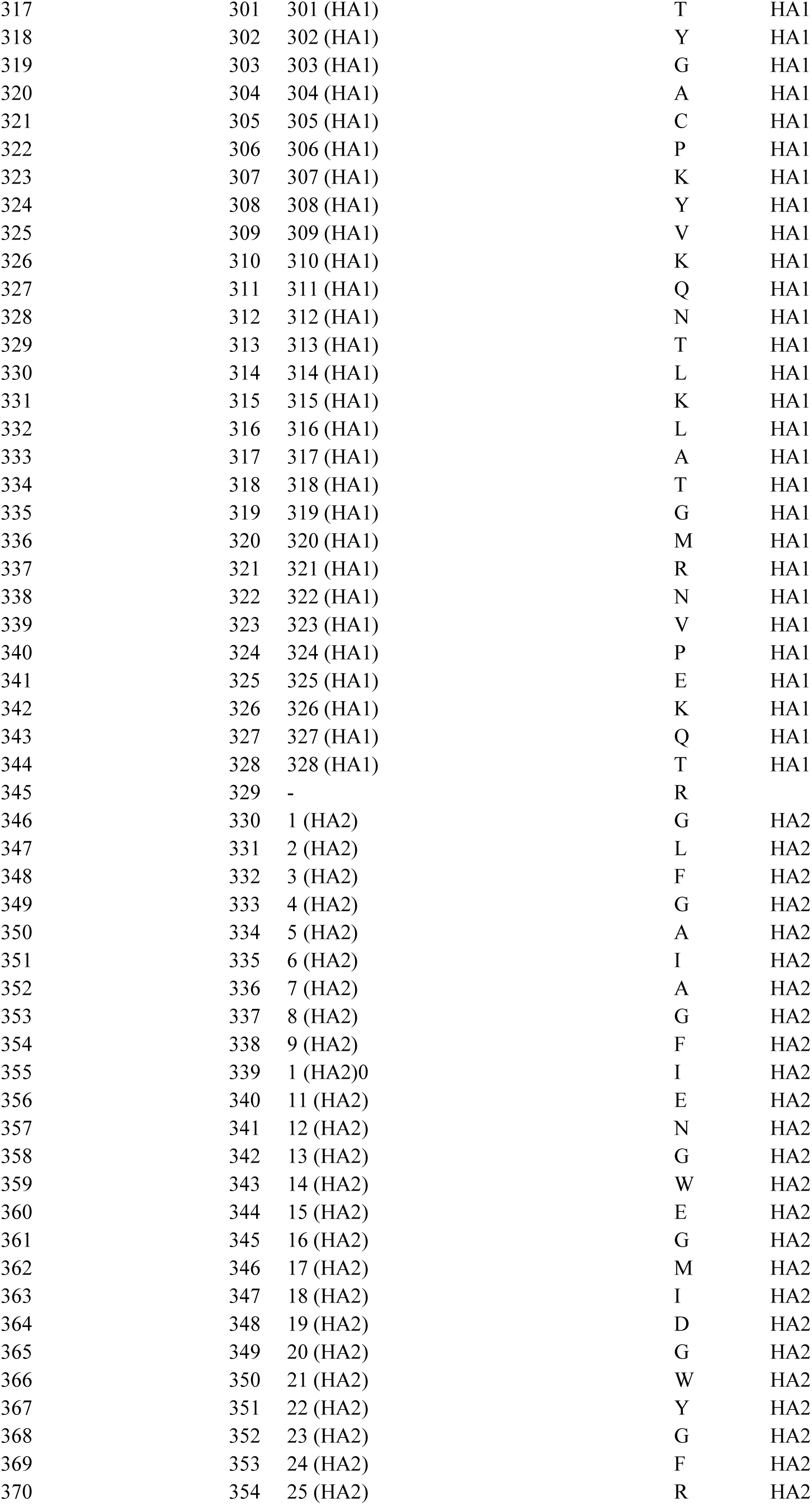
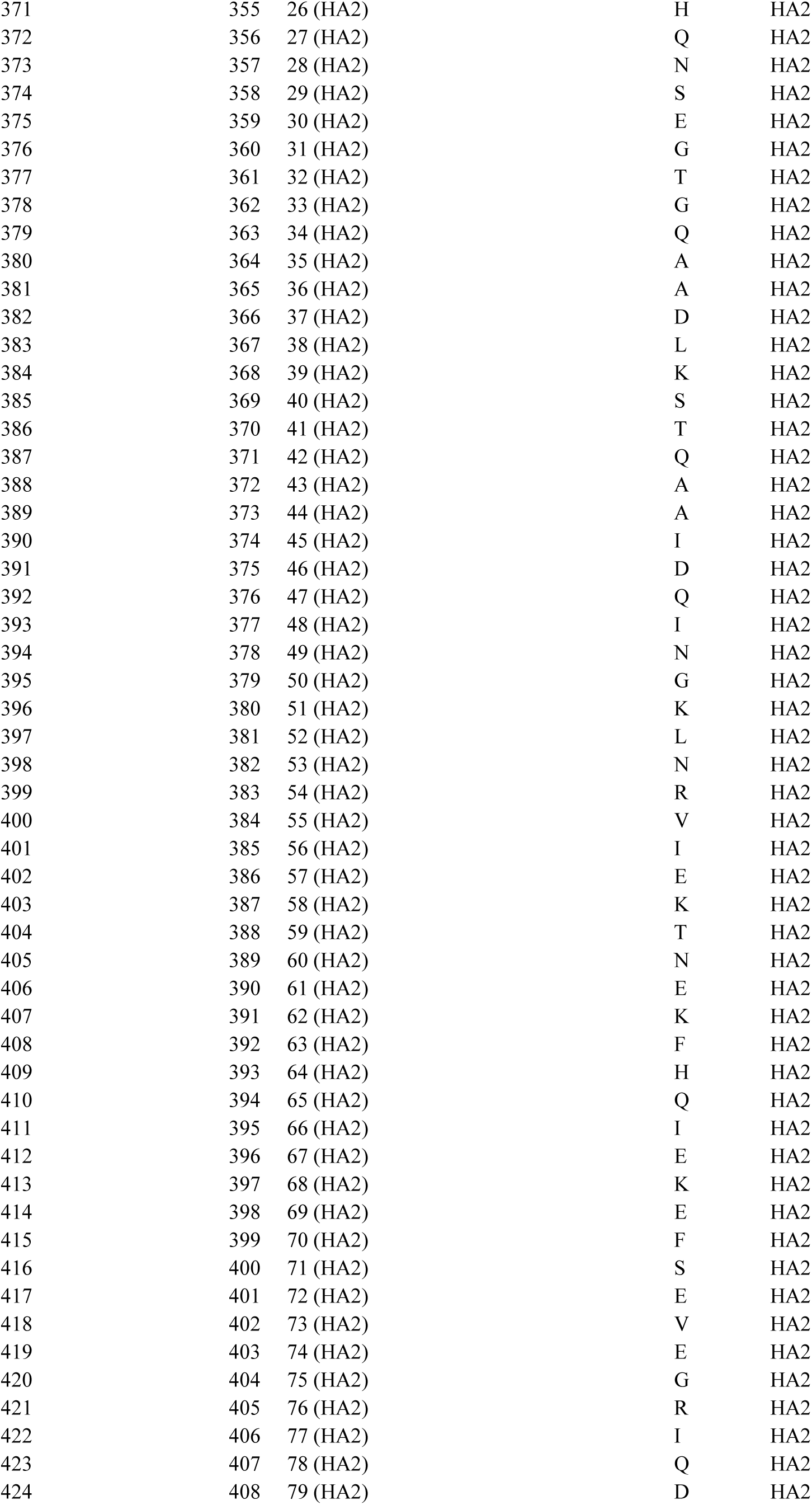
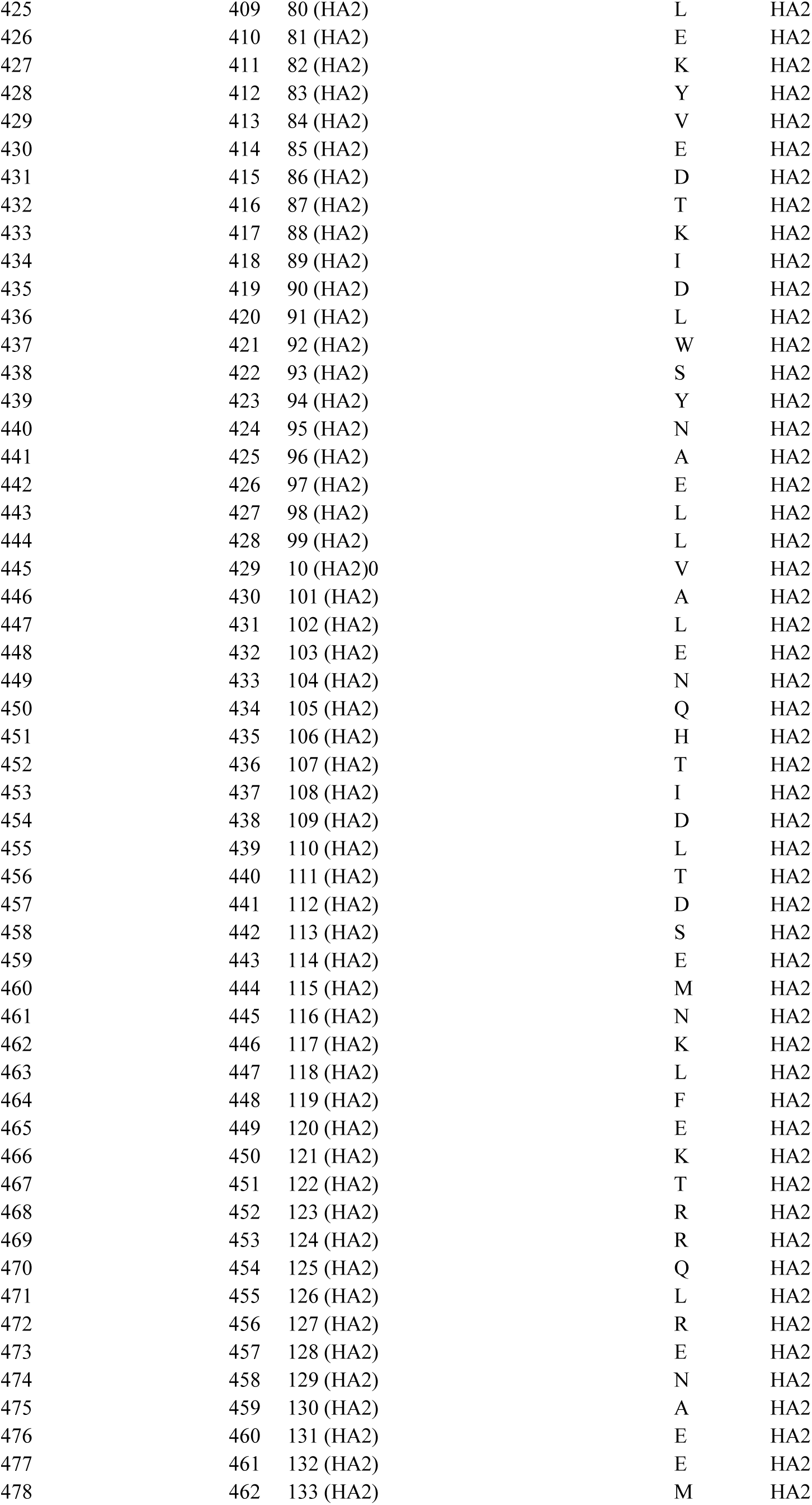
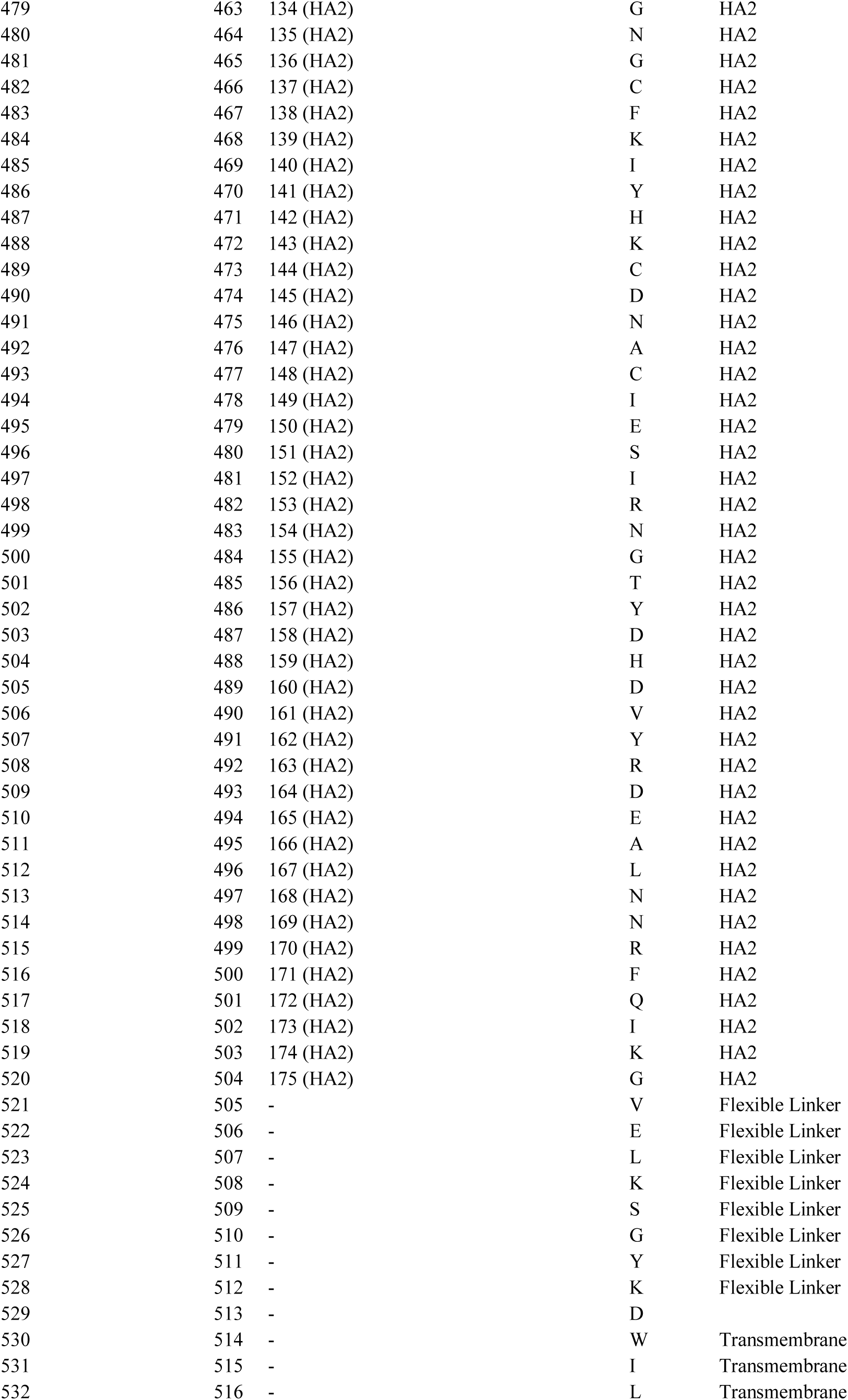
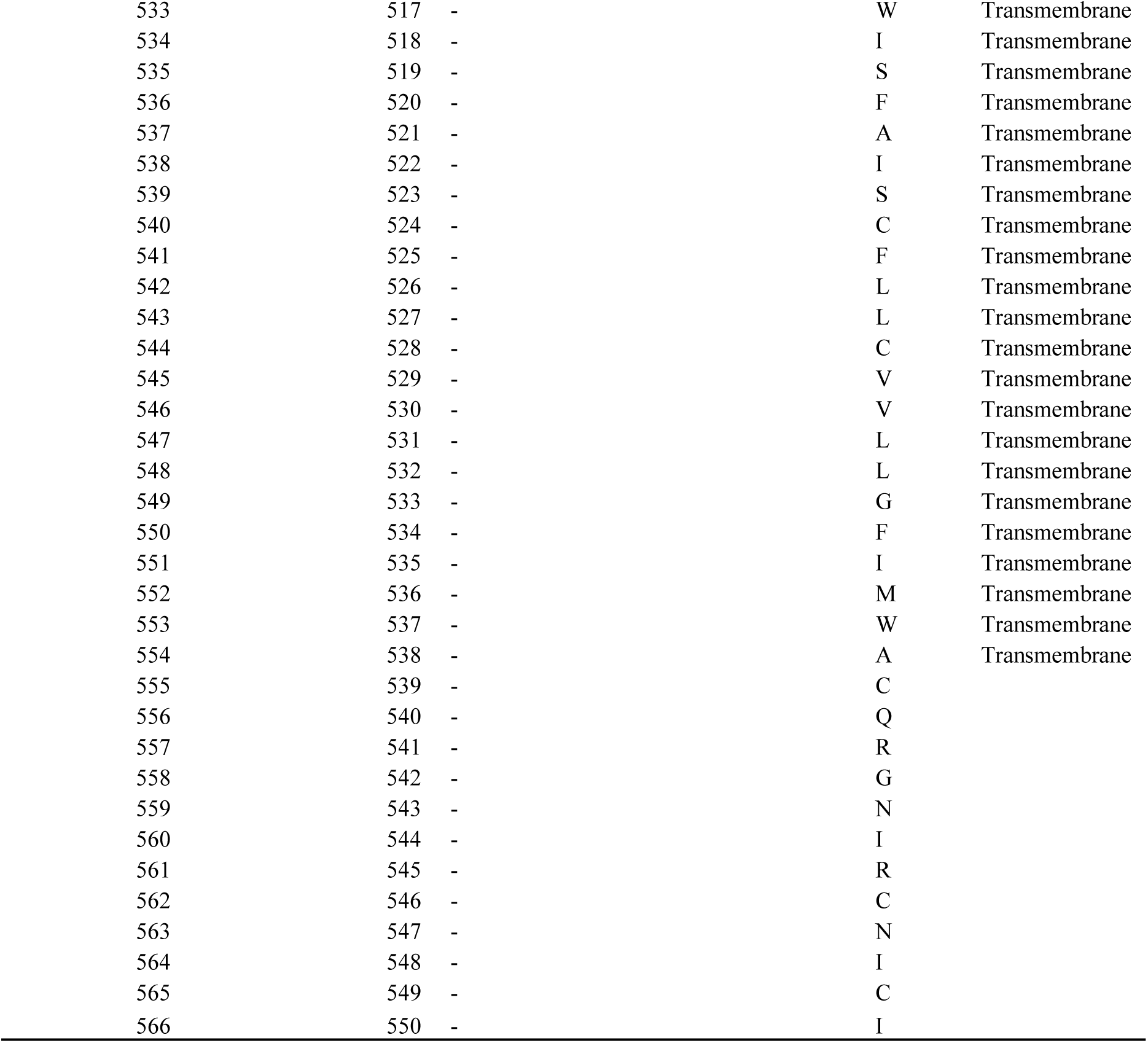
Comparison of multiple HA numbering schemes using a H3 (A/Aichi/2/1968(H3N2)) template. Template for FLU DB numbering is H3. Flexible linker and transmembrane domain were located by aligning to A/duck/Alberta/35/76(H1N1), PDB ID: 6HJR.

## References

1. Koel, B.F. et al. Substitutions near the receptor binding site determine major antigenic change during influenza virus evolution. Science 342, 976–979 (2013).

2. Sun, W. et al. Development of influenza B universal vaccine candidates using the “Mosaic” hemagglutinin approach. Journal of virology 93 (2019).

3. Krammer, F. & Palese, P. Universal influenza virus vaccines that target the conserved hemagglutinin stalk and conserved sites in the head domain. The Journal of infectious diseases 219, S62–S67 (2019).

4. Carter, D.M. et al. Design and characterization of a computationally optimized broadly reactive hemagglutinin vaccine for H1N1 influenza viruses. Journal of virology 90, 4720–4734 (2016).

5. Gibson, D.G. et al. Enzymatic assembly of DNA molecules up to several hundred kilobases. Nature methods 6, 343–345 (2009).

6. Burney, L.E. Influenza immunization: statement. Public health reports 75, 944 (1960).

7. Flannery, B., et al. Interim estimates of 2017–18 seasonal influenza vaccine effectiveness—United States, February 2018. Morbidity and Mortality Weekly Report 67, 180 (2018).

8. Xie, H. et al. H3N2 mismatch of 2014–15 northern hemisphere influenza vaccines and head-to-head comparison between human and ferret antisera derived antigenic maps. Scientific reports 5, 1–10 (2015).

9. Chiu, C. et al. Cross-reactive humoral responses to influenza and their implications for a universal vaccine. Annals of the New York Academy of Sciences 1283, 13–21 (2013).

10. Hagan, T. et al. Antibiotics-driven gut microbiome perturbation alters immunity to vaccines in humans. Cell 178, 1313–1328. e1313 (2019).

11. Henry, C. et al. Monoclonal Antibody Responses after Recombinant Hemagglutinin Vaccine versus Subunit Inactivated Influenza Virus Vaccine: a Comparative Study. Journal of Virology 93, e01150–01119 (2019).

12. Staneková, Z. & Varečková, E. Conserved epitopes of influenza A virus inducing protective immunity and their prospects for universal vaccine development. Virology journal 7, 1–13 (2010).

13. Burke, D.F. & Smith, D.J. A recommended numbering scheme for influenza A HA subtypes. PloS one 9 (2014).

14. Kirkpatrick, E., Qiu, X., Wilson, P.C., Bahl, J. & Krammer, F. The influenza virus hemagglutinin head evolves faster than the stalk domain. Scientific reports 8, 1–14 (2018).

15. Deem, M.W. & Pan, K. The epitope regions of H1-subtype influenza A, with application to vaccine efficacy. Protein Engineering, Design & Selection 22, 543–546 (2009).

16. Hai, R. et al. Influenza viruses expressing chimeric hemagglutinins: globular head and stalk domains derived from different subtypes. Journal of virology 86, 5774–5781 (2012).

17. Steel, J. et al. Influenza virus vaccine based on the conserved hemagglutinin stalk domain. MBio 1 (2010).

18. Medina, R.A. et al. Glycosylations in the globular head of the hemagglutinin protein modulate the virulence and antigenic properties of the H1N1 influenza viruses. Science translational medicine 5, 187ra170–187ra170 (2013).

19. Li, L. et al. Multi-task learning sparse group lasso: a method for quantifying antigenicity of influenza A (H1N1) virus using mutations and variations in glycosylation of Hemagglutinin. BMC bioinformatics 21, 1–22 (2020).

20. Crooks, G.E., Hon, G., Chandonia, J.-M. & Brenner, S.E. WebLogo: a sequence logo generator. Genome research 14, 1188–1190 (2004).

21. Schrödinger, L. The PyMOL Molecular Graphics System, Version 2.0 Schrödinger, LLC (2017). Google Scholar There is no corresponding record for this reference.

22. Pettersen, E.F. et al. UCSF Chimera—a visualization system for exploratory research and analysis. Journal of computational chemistry 25, 1605–1612 (2004).

23. Abola, E.E., Bernstein, F.C. & Koetzle, T.F. in Neutrons in Biology 441–441 (Springer, 1984).

24. Weis, W.I., Brunger, A.T., Skehel, J.J. & Wiley, D.C. Refinement of the influenza virus hemagglutinin by simulated annealing. J Mol Biol 212, 737–761 (1990).

25. Zhang, W. et al. Molecular basis of the receptor binding specificity switch of the hemagglutinins from both the 1918 and 2009 pandemic influenza A viruses by a D225G substitution. J Virol 87, 5949–5958 (2013).

26. McDonald, N.J., Smith, C.B. & Cox, N.J. Antigenic drift in the evolution of H1N1 influenza A viruses resulting from deletion of a single amino acid in the haemagglutinin gene. Journal of General Virology 88, 3209–3213 (2007).

27. Benton, D.J. et al. Influenza hemagglutinin membrane anchor. Proceedings of the National Academy of Sciences 115, 10112–10117 (2018).

28. Whittle, J.R. et al. Flow cytometry reveals that H5N1 vaccination elicits cross-reactive stem-directed antibodies from multiple Ig heavy-chain lineages. J Virol 88, 4047–4057 (2014).

29. Setliff, I. et al. High-Throughput Mapping of B Cell Receptor Sequences to Antigen Specificity. Cell 179, 1636–1646 e1615 (2019).

30. Harvey, W.T. et al. Identification of low-and high-impact hemagglutinin amino acid substitutions that drive antigenic drift of influenza A (H1N1) viruses. PLoS pathogens 12 (2016).

31. Li, L., DeLiberto, T.J., Killian, M.L., Torchetti, M.K. & Wan, X.-F. Evolutionary pathway for the 2017 emergence of a novel highly pathogenic avian influenza A (H7N9) virus among domestic poultry in Tennessee, United States. Virology 525, 32–39 (2018).

32. Krammer, F., Li, L. & Wilson, P.C. Emerging from the Shadow of Hemagglutinin: Neuraminidase Is an Important Target for Influenza Vaccination. Cell Host & Microbe 26, 712–713 (2019).

33. Zhu, X. et al. Structural basis of protection against H7N9 influenza virus by human anti-N9 neuraminidase antibodies. Cell host & microbe 26, 729–738. e724 (2019).

34. Gilchuk, I.M. et al. Influenza H7N9 virus neuraminidase-specific human monoclonal antibodies inhibit viral egress and protect from lethal influenza infection in mice. Cell host & microbe 26, 715–728. e718 (2019).

35. Chen, Y.-Q. et al. Influenza infection in humans induces broadly cross-reactive and protective neuraminidase-reactive antibodies. Cell 173, 417–429. e410 (2018).

36. Bao, Y. et al. The influenza virus resource at the National Center for Biotechnology Information. Journal of virology 82, 596–601 (2008).

37. Bogner, P., Capua, I., Lipman, D.J. & Cox, N.J. A global initiative on sharing avian flu data. Nature 442, 981–981 (2006).

38. Edgar, R.C. MUSCLE: multiple sequence alignment with high accuracy and high throughput. Nucleic acids research 32, 1792–1797 (2004).

39. Sievers, F. et al. Fast, scalable generation of high-quality protein multiple sequence alignments using Clustal Omega. Molecular systems biology 7 (2011).

40. Stamatakis, A. RAxML version 8: a tool for phylogenetic analysis and post-analysis of large phylogenies. Bioinformatics 30, 1312–1313 (2014).

