## SupplementaryData for "Librator, a platform for optimized sequence editing, design, and expression of influenza virus proteins"

**Dataset S1**

Single mutants between HK-14 and SWZ-13

***Recipes***

| **Strain name** | **Fragment1** | **Fragment2** | **Fragment3** | **Fragment4** |
| --- | --- | --- | --- | --- |
| A/Hong Kong/4801/2014 | H3-F1-0001 | H3-F2-0001 | H3-F3-0001 | H3-F4-0001 |
| A/Switzerland/9715293/2013-G158R(Cocktail) | H3-F1-0002 | H3-F2-0002 | H3-F3-0002 | H3-F4-0002 |
| A/Switzerland/9715293/2013-S154A(Cocktail) | H3-F1-0002 | H3-F2-0003 | H3-F3-0002 | H3-F4-0002 |
| A/Switzerland/9715293/2013-R342K(Cocktail) | H3-F1-0002 | H3-F2-0004 | H3-F3-0003 | H3-F4-0002 |
| A/Switzerland/9715293/2013-L19I(Cocktail) | H3-F1-0001 | H3-F2-0004 | H3-F3-0002 | H3-F4-0002 |
| A/Switzerland/9715293/2013-D505N(Cocktail) | H3-F1-0002 | H3-F2-0004 | H3-F3-0002 | H3-F4-0001 |
| A/Switzerland/9715293/2013-A144T(Cocktail) | H3-F1-0002 | H3-F2-0005 | H3-F3-0002 | H3-F4-0002 |
| A/Switzerland/9715293/2013-K176T(Cocktail) | H3-F1-0002 | H3-F2-0006 | H3-F3-0002 | H3-F4-0002 |
| A/Switzerland/9715293/2013-N160S(Cocktail) | H3-F1-0002 | H3-F2-0007 | H3-F3-0002 | H3-F4-0002 |
| A/Switzerland/9715293/2013-Q327H(Cocktail) | H3-F1-0002 | H3-F2-0004 | H3-F3-0004 | H3-F4-0002 |
| A/Switzerland/9715293/2013-S175Y(Cocktail) | H3-F1-0002 | H3-F2-0008 | H3-F3-0002 | H3-F4-0002 |
| A/Switzerland/9715293/2013 | H3-F1-0002 | H3-F2-0004 | H3-F3-0002 | H3-F4-0002 |

***Fragments***

H3-F1-0001, H3-F1-0002, H3-F2-0001, H3-F2-0002, H3-F2-0003, H3-F2-0004, H3-F2-0005, H3-F2-0006, H3-F2-0007, H3-F2-0008, H3-F3-0001, H3-F3-0002, H3-F3-0003, H3-F3-0004, H3-F4-0001, H3-F4-0002

***Full HA cost***

| number | unit price | total price |
| --- | --- | --- |
| 12 | 229 | 2748 |

***Librator fragment cost***

| number | unit price | total price | save cost | save pct |
| --- | --- | --- | --- | --- |
| 16 | 79 | 1264 | 1484 | 54% |

**Dataset S2**

39 H3 sequences from 1968 to 2018.

***Recipes***

| **Strain name** | **Fragment1** | **Fragment2** | **Fragment3** | **Fragment4** |
| --- | --- | --- | --- | --- |
| A/Hong Kong/1/1968 | H3-F1-0001 | H3-F2-0001 | H3-F3-0001 | H3-F4-0001 |
| A/Victoria/3/1975 | H3-F1-0002 | H3-F2-0002 | H3-F3-0002 | H3-F4-0002 |
| A/Bilthoven/9459/1974 | H3-F1-0003 | H3-F2-0003 | H3-F3-0003 | H3-F4-0003 |
| A/England/42/1972 | H3-F1-0004 | H3-F2-0004 | H3-F3-0004 | H3-F4-0004 |
| A/Port Chalmers/1/1973 | H3-F1-0005 | H3-F2-0005 | H3-F3-0005 | H3-F4-0002 |
| A/Fujian/411/2002 | H3-F1-0006 | H3-F2-0006 | H3-F3-0006 | H3-F4-0005 |
| A/California/7/2004 | H3-F1-0006 | H3-F2-0007 | H3-F3-0007 | H3-F4-0006 |
| A/Wisconsin/67/2005 | H3-F1-0007 | H3-F2-0008 | H3-F3-0008 | H3-F4-0007 |
| A/Brisbane/10/2007 | H3-F1-0008 | H3-F2-0009 | H3-F3-0009 | H3-F4-0008 |
| A/TayNguyen/TN410/2005 | H3-F1-0006 | H3-F2-0010 | H3-F3-0010 | H3-F4-0009 |
| A/Victoria/361/2011 | H3-F1-0009 | H3-F2-0011 | H3-F3-0011 | H3-F4-0008 |
| A/Egypt/N08920/2009 | H3-F1-0010 | H3-F2-0012 | H3-F3-0012 | H3-F4-0008 |
| A/Perth/16/2009_A147E_HA2 | H3-F1-0011 | H3-F2-0013 | H3-F3-0012 | H3-F4-0010 |
| A/Perth/16/2009_E325S_HA1 | H3-F1-0011 | H3-F2-0013 | H3-F3-0013 | H3-F4-0008 |
| A/Perth/16/2009_L38Q_HA2 | H3-F1-0011 | H3-F2-0013 | H3-F3-0014 | H3-F4-0008 |
| A/Perth/16/2009_N53G_HA2 | H3-F1-0011 | H3-F2-0013 | H3-F3-0015 | H3-F4-0008 |
| A/Perth/16/2009_S29Y_HA2 | H3-F1-0011 | H3-F2-0013 | H3-F3-0016 | H3-F4-0008 |
| A/Perth/16/2009 | H3-F1-0011 | H3-F2-0013 | H3-F3-0012 | H3-F4-0008 |
| A/Hong Kong/4801/2014 | H3-F1-0012 | H3-F2-0014 | H3-F3-0017 | H3-F4-0011 |
| A/Switzerland/8060/2017 | H3-F1-0013 | H3-F2-0015 | H3-F3-0018 | H3-F4-0011 |
| A/England/80740425/2018 | H3-F1-0014 | H3-F2-0016 | H3-F3-0019 | H3-F4-0012 |
| A/Singapore/INFIMH-16-0019/2016 | H3-F1-0012 | H3-F2-0017 | H3-F3-0017 | H3-F4-0013 |
| A/Kansas/14/2017 | H3-F1-0015 | H3-F2-0018 | H3-F3-0020 | H3-F4-0014 |
| A/Switzerland/9715293/2013 | H3-F1-0016 | H3-F2-0019 | H3-F3-0020 | H3-F4-0008 |
| A/Bangkok/1/1979 | H3-F1-0017 | H3-F2-0020 | H3-F3-0021 | H3-F4-0015 |
| A/Texas/1/1977 | H3-F1-0018 | H3-F2-0021 | H3-F3-0022 | H3-F4-0016 |
| A/New York/680/1995 | H3-F1-0019 | H3-F2-0022 | H3-F3-0023 | H3-F4-0017 |
| A/Beijing/353/1989 | H3-F1-0020 | H3-F2-0023 | H3-F3-0024 | H3-F4-0018 |
| A/Beijing/32/1992 | H3-F1-0020 | H3-F2-0024 | H3-F3-0025 | H3-F4-0018 |
| A/Shangdong/9/1993 | H3-F1-0021 | H3-F2-0025 | H3-F3-0026 | H3-F4-0018 |
| A/Memphis/2/1986 | H3-F1-0022 | H3-F2-0026 | H3-F3-0027 | H3-F4-0019 |
| A/Colorado/2/1986 | H3-F1-0023 | H3-F2-0027 | H3-F3-0028 | H3-F4-0018 |
| A/Sichuan/2/1987 | H3-F1-0024 | H3-F2-0028 | H3-F3-0029 | H3-F4-0018 |
| A/Auckland/610/2002 | H3-F1-0025 | H3-F2-0029 | H3-F3-0030 | H3-F4-0020 |
| A/Panama/2007/1999 | H3-F1-0026 | H3-F2-0030 | H3-F3-0031 | H3-F4-0021 |
| A/Sydney/5/1997 | H3-F1-0027 | H3-F2-0031 | H3-F3-0031 | H3-F4-0022 |
| A/Hong Kong/358/1996 | H3-F1-0028 | H3-F2-0032 | H3-F3-0032 | H3-F4-0017 |
| A/Paris/908/97 | H3-F1-0029 | H3-F2-0033 | H3-F3-0032 | H3-F4-0017 |
| A/Wuhan/359/1995 | H3-F1-0030 | H3-F2-0034 | H3-F3-0032 | H3-F4-0017 |

***Fragments***

H3-F1-0001, H3-F1-0002, H3-F1-0003, H3-F1-0004, H3-F1-0005, H3-F1-0006, H3-F1-0007, H3-F1-0008, H3-F1-0009, H3-F1-0010, H3-F1-0011, H3-F1-0012, H3-F1-0013, H3-F1-0014, H3-F1-0015, H3-F1-0016, H3-F1-0017, H3-F1-0018, H3-F1-0019, H3-F1-0020, H3-F1-0021, H3-F1-0022, H3-F1-0023, H3-F1-0024, H3-F1-0025, H3-F1-0026, H3-F1-0027, H3-F1-0028, H3-F1-0029, H3-F1-0030, H3-F2-0001, H3-F2-0002, H3-F2-0003, H3-F2-0004, H3-F2-0005, H3-F2-0006, H3-F2-0007, H3-F2-0008, H3-F2-0009, H3-F2-0010, H3-F2-0011, H3-F2-0012, H3-F2-0013, H3-F2-0014, H3-F2-0015, H3-F2-0016, H3-F2-0017, H3-F2-0018, H3-F2-0019, H3-F2-0020, H3-F2-0021, H3-F2-0022, H3-F2-0023, H3-F2-0024, H3-F2-0025, H3-F2-0026, H3-F2-0027, H3-F2-0028, H3-F2-0029, H3-F2-0030, H3-F2-0031, H3-F2-0032, H3-F2-0033, H3-F2-0034, H3-F3-0001, H3-F3-0002, H3-F3-0003, H3-F3-0004, H3-F3-0005, H3-F3-0006, H3-F3-0007, H3-F3-0008, H3-F3-0009, H3-F3-0010, H3-F3-0011, H3-F3-0012, H3-F3-0013, H3-F3-0014, H3-F3-0015, H3-F3-0016, H3-F3-0017, H3-F3-0018, H3-F3-0019, H3-F3-0020, H3-F3-0021, H3-F3-0022, H3-F3-0023, H3-F3-0024, H3-F3-0025, H3-F3-0026, H3-F3-0027, H3-F3-0028, H3-F3-0029, H3-F3-0030, H3-F3-0031, H3-F3-0032, H3-F4-0001, H3-F4-0002, H3-F4-0003, H3-F4-0004, H3-F4-0005, H3-F4-0006, H3-F4-0007, H3-F4-0008, H3-F4-0009, H3-F4-0010, H3-F4-0011, H3-F4-0012, H3-F4-0013, H3-F4-0014, H3-F4-0015, H3-F4-0016, H3-F4-0017, H3-F4-0018, H3-F4-0019, H3-F4-0020, H3-F4-0021, H3-F4-0022

***Full HA cost***

| number | unit price | total price |
| --- | --- | --- |
| 39 | 229 | 8931 |

***Librator fragment cost***

| number | unit price | total price | save cost | save pct |
| --- | --- | --- | --- | --- |
| 118 | 79 | 9322 | -391 | -4% |
